# P-cadherin mechanoactivates tumor–mesothelium metabolic coupling to promote ovarian cancer metastasis

**DOI:** 10.1101/2024.06.02.597059

**Authors:** Jing Ma, Sally K. Y. To, Katie S. W. Fung, Kun Wang, Jiangwen Zhang, Alfonso H. W. Ngan, Susan Yung, Tak-Mao Chan, Carmen C. L. Wong, Philip P. C. Ip, Ling Peng, Hong-Yan Guo, Chi Bun Chan, Alice S.T. Wong

**Affiliations:** School of Biological Sciences, University of Hong Kong, Pokfulam Road, Hong Kong; Department of Pharmacy, South China Hospital, Medical School, Shenzhen University, Shenzhen, 518116, China; Laboratory for Synthetic Chemistry and Chemical Biology Limited, 17W, Hong Kong Science and Technology Parks, New Territories, Hong Kong; Department of Mechanical Engineering, University of Hong Kong, Pokfulam Road, Hong Kong; Department of Medicine, School of Clinical Medicine, University of Hong Kong, Queen Mary Hospital, Sassoon Road, Hong Kong; Department of Pathology, School of Clinical Medicine, University of Hong Kong, Queen Mary Hospital, Sassoon Road, Hong Kong; Aix-Marseille Université, CNRS, Centre Interdisciplinaire de Nanoscience de Marseille, UMR 13288 Marseille, France; Department of Obstetrics and Gynecology, Peking University Third Hospital, Beijing 100191, China

**Author notes:** Correspondence: Alice Sze Tsai Wong,; Chi Bun Chan. These authors made equal contributions (Jing Ma, Sally K. Y. To, Katie S. W. Fung, Kun Wang).

**Keywords:** P-cadherin, tumor-mesothelium, mechanotransduction, lipogenesis, glycolysis

## Abstract

Peritoneal metastasis exacerbates the prognosis of ovarian cancer patients. Adhesion of cancer cells to mesothelium is a rate-limiting prerequisite for this process. How metastatic cells sense and respond to the dynamic biomechanical microenvironment at the mesothelial niche to initiate metastatic lesions remains unclear. Here, the study demonstrates that highly metastatic (HM), but not non-metastatic (NM) ovarian cancer cells, selectively activate the peritoneal mesothelium. Atomic force microscopy reveals that HM cells exert increased adhesive force on mesothelial cells via P-cadherin, a cell-cell adhesion molecule abundant in late-stage tumors. Transcriptomic and molecular analyses show that mechanical induction of P-cadherin enhances lipogenic gene expression and lipid content in HM cells by SREBP1. P-cadherin activation does not affect lipogenic activity but induces glycolysis in the interacting mesothelium. Targeted lipidomic analysis reveals that lactate produced by the glycolytic mesothelium facilitates metastatic outgrowth as a direct substrate for *de novo* lipogenesis. Inhibiting lactate shuttling via nanodelivery of siRNA targeting P-cadherin or MCT1/4 transporters significantly suppresses metastasis in mice. The association of high fatty acid synthase in patient metastatic samples and increased P-cadherin expression supports enhanced *de novo* lipogenesis in the metastatic niche. The study reveals P-cadherin-mediated mechano-metabolic coupling as a promising target to restrain peritoneal metastasis.

## Main

Ovarian cancer is the leading cause of mortality among gynecological malignancies. Metastatic ovarian cancer cells exhibit a marked predilection for colonizing the peritoneum and omentum^1,2^. Peritoneal metastasis, although less frequent than blood-borne metastases, poses significant challenges for treatment due to its rapid and expansive dissemination fueled by positive feedback mechanisms. The aggressive nature of this dissemination mode renders current therapies largely ineffective, resulting in a 5-year survival rate of less than 25% for advanced-stage cases^3^.

Metastasis is a complex and highly inefficient process, with less than 0.01% of disseminated cancer cells successfully establishing themselves in a target tissue and initiating new metastatic growth^4^. In the context of peritoneal metastasis, adhesion to the mesothelium lining at the peritoneal surface is a first critical step^5^. Mesothelial cells were traditionally viewed as passive barriers, but emerging research challenges this perception and suggests their active involvement in shaping the metastatic microenvironment in ovarian cancer. Recent studies have revealed that cancer-associated mesothelial cells could secrete extracellular matrix proteins such as fibronectin and collagen, cytokines and chemokines like interleukin-8 and CCL2, as well as soluble proteins like angiopoietin-like 4 and osteopontin, which may collectively create a favorable environment that promotes tumor adhesion, chemoresistance and also immune evasion^6,7,8,9,10,11^. On the other hand, metastatic cancer cells have been found to induce mesenchymal transition in mesothelial cells via interleukin-1 and transforming growth factor β^8,11^. However, these studies have primarily focused on secreted factors, the involvement of cell-cell contact in signal transduction remains largely unknown.

Metastatic cells possess remarkable capability to sense and respond to a wide range of biochemical and biophysical signals originating from their surrounding microenvironment. While the biomechanics of tumor microenvironment, known as the mechano-niche, have long been overlooked, they are now recognized as distinct features in cancer development and progression^12,13^. Mechanosensitive proteins have emerged as promising and unique targets for cancer therapeutics. The mechano-niche encompasses various factors, such as tissue stiffness, extracellular matrix composition, and mechanical forces, which can profoundly influence the behaviors and functions of cancer cells. In addition to the dynamic mechanical changes, metastatic cells must also adapt to the diverse metabolic demands encountered during their journey. Significant rewiring of metabolic circuits, including aerobic glycolysis, oxidative phosphorylation, lipid metabolism, and glutamine metabolism, is regarded as a hallmark of cancer. However, the precise mechanisms through which the mechano-niche impacts cancer cell metabolism remain largely unknown.

Cadherins are well-known mechanosensitive sensors that transmit both outside–in and inside–out signals which modulate their conformation and adhesive properties^14,15^. E-cadherin-mediated mechanotransduction was shown to regulate glucose uptake through the activation of adenosine monophosphate-activated protein kinase^16^, while another study reported that heterophilic E-cadherin/N-cadherin mechanical signaling could drive collective migration^17^. Despite these findings, the functional significance and molecular mechanisms of mechanical signaling by cadherins are largely unexplored. Our present study aims to investigate the potential role of cadherin in establishing a connection between cell adhesion mechanics and metabolic reprogramming at the mesothelial metastatic niche. Our results reveal a unique involvement of P-cadherin, distinguishing it from other cadherins, in activating the peritoneal niche and establishing metabolic coupling with the mesothelium during peritoneal colonization. These significant findings underscore the therapeutic potential of targeting P-cadherin-mediated mechanotransduction for the treatment of peritoneal metastasis.

## Results

### P-cadherin promotes the adhesion of metastatic cells to the mesothelium

To elucidate the molecular basis of metastasis under tightly controlled conditions, we have established an isogenic model of spontaneous human ovarian cancer metastasis consisting of the highly metastatic (HM)-non-metastatic (NM) cell pair. Orthotopic injection in both immunocompromised and humanized mouse models reveled that HM cells rapidly metastasized within the peritoneum, closely mirroring human ovarian cancer dissemination patterns, whereas NM cells did not metastasize despite similar tumorigenicity as HM cells^18,19^. To ensure that our observations were not cell-line specific, we have used HM-NM pairs derived from HEYA8 and OVSAHO cell lines. To mimic tumor-mesothelial interaction, we conducted mesothelial adhesion assays, wherein tumor cells were allowed to adhere to a monolayer of mesothelial cells, followed by gentle washing steps to remove non-adherent cells. Both primary human peritoneal mesothelial cells (HPMC) and human mesothelial cell line MeT5A were employed to ensure the robustness of our observations. A significant increase in the adhesion of HEYA8 HM cells to mesothelial cell monolayer compared to their NM counterparts was observed (HPMC: 3.40 ± 0.25 fold increase, MeT5A: 7.14 ± 0.64-fold increase; Fig. 1a). Similar results were obtained when using the OVSAHO HM–NM cell pair (4.54 ± 0.40-fold increase; Supplementary Fig. 1a), indicating that metastatic cells are more capable of adhering to the mesothelium. Conditioned medium from either HM or MeT5A cells alone did not affect HM–MeT5A adhesion (Supplementary Fig. 1b), further suggesting the involvement of a contact-dependent mechanism in this tumor–mesothelium interaction.

**Fig. 1.**
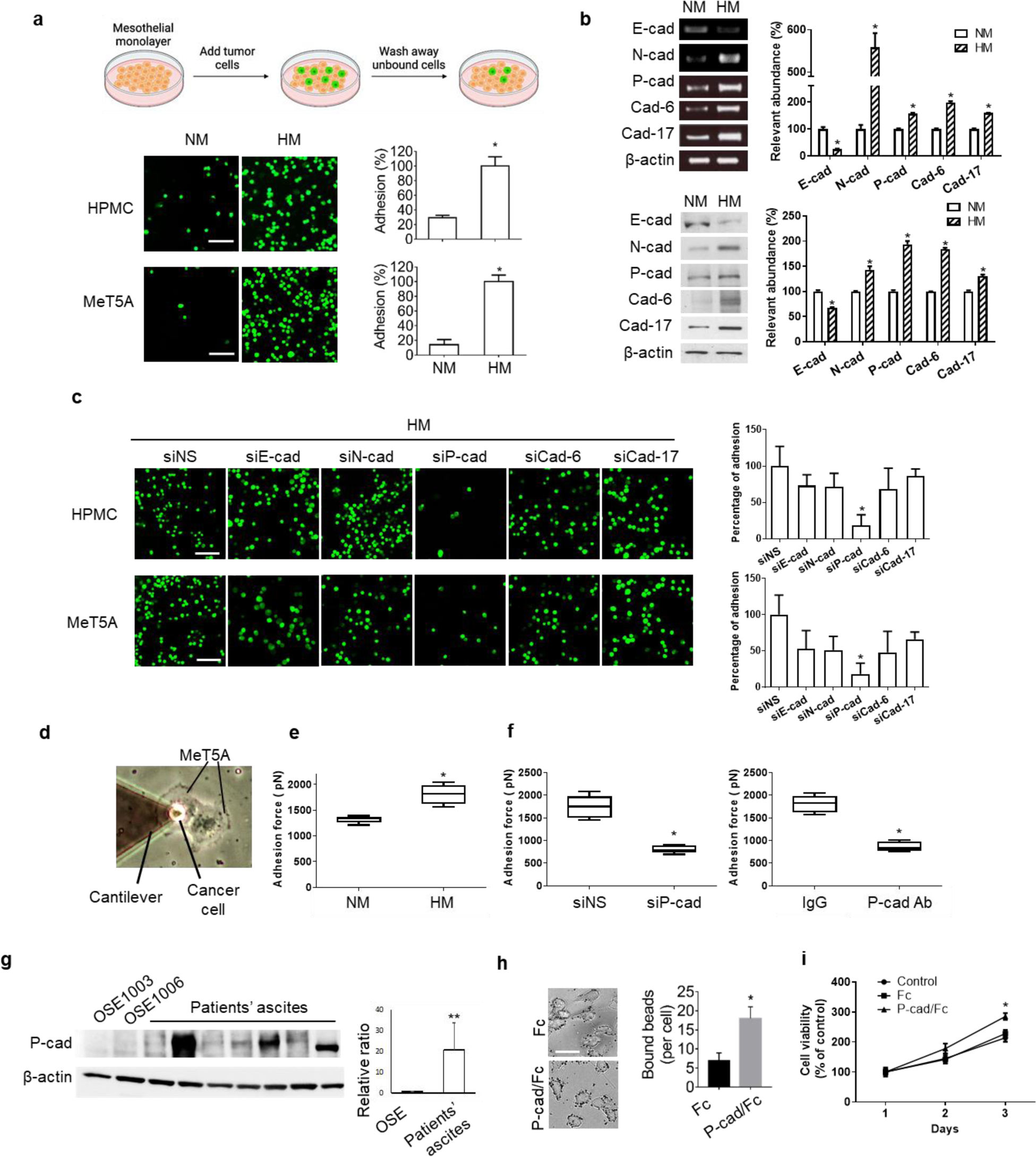
P-cadherin mechanically modulates tumor-mesothelial adhesion. **a,** Non-metastatic (NM) and highly metastatic (HM) derived from HEYA8 ovarian cancer cells were allowed to adhere onto confluent human peritoneal mesothelial cell (HPMC) or MeT5A monolayer, followed by gentle washing to remove unbound cells. Representative fluorescent images and quantification of adhered cells were shown. Scale bar, 100 μm. **b,** Expression levels of P-cadherin (P-cad), E-cadherin (E-cad), N-cadherin (N-cad), cadherin-6 (Cad-6), and cadherin-17 (Cad-17) in NM and HM cells were analyzed by RT-PCR and Western blot analysis, with β-actin as the loading control. **c,** HM cells transfected with non-specific, E-cad, N-cad, P-cad, Cad-6 or Cad-17 siRNA (siNS, siE-cad, siN-cad, siP-cad, siCad-6, siCad-17, respectively) were allowed to adhere onto HPMC or MeT5A, followed by gentle washing to remove unbound cells. Representative fluorescent images were shown. At least five microscopic fields of the adhered cells were quantified for each condition and data are presented as mean ± SD. Scale bar, 100 μm. **d,** An image illustrating the measurement of cell-cell rupture forces by atomic force microscopy was shown. **e,** Rupture forces of NM or HM detached from MeT5A cells were measured (n=5). **f,** Rupture forces of HM detached from MeT5A cells in the absence or presence of siP-cad or P-cadherin blocking antibody (P-cad Ab) were measured, with siNS and IgG as controls respectively (n=5). **g,** Protein expression of P-cad in normal ovarian surface epithelial (OSE) cells and ovarian caner patient ascites (n=7) were analyzed by Western blot. β-actin serves as the loading control. **h,** HM cells were incubated with Fc- or P-cad/Fc-coated beads for 30 min. Bound beads were quantified after washing with PBS for three times. Scale bar, 20 μm. **i,** Cell viability of HM cells treated with Fc or P-cad/Fc coated beads for 3 days were determined. All experiments were repeated three times. Data are presented as mean ± SD. \**P* < 0.05, *\*\* P* < 0.01.

To identify the cell adhesion molecule responsible for mediating the interaction between HM cells and the mesothelium, we examined the expression of E-cadherin, N-cadherin, P-cadherin, cadherin-6, and cadherin-17, which have been previously implicated in the progression of ovarian cancer^20,21,22^. Our analysis, at both the mRNA and protein levels, showed significantly lower expression of E-cadherin and significantly higher expression of the other cadherins in HEYA8 HM cells compared to NM cells (Fig. 1b). Similar protein expression changes were observed in OVSAHO cell pair (Supplementary Fig. 1c). To identify which of these cadherins is important for the interaction, we used corresponding small interfering RNAs (siRNAs) to target E-cadherin (siE-cad), N-cadherin (siN-cad), P-cadherin (siP-cad), cadherin-6 (siCad-6), and cadherin-17 (siCad-17). We found that specifically blocking P-cadherin resulted in a significant decrease in the mesothelial adhesion of HEYA8 HM cells. The adhesion of HEYA8 HM to HPMC or MeT5A decreased notably with siP-cad (83 ± 7%), whereas siE-cad (27 ± 15%), siN-cad (29 ± 19%), siCad-6 (36 ± 9%), and siCad-17 (10 ± 10%) did not exhibit significant reductions compared to a non-specific siRNA (siNS) control (Fig. 1c). All siRNA showed similar knockdown efficiency (Supplementary Fig. 1d). Additionally, siP-cad significantly reduced adhesion of OVSAHO HM cells (HPMC: 68 ± 11% and MeT5A: 77 ± 8%) (Supplementary Fig. 1e). The expression of P-cadherin was detected in both HPMC and MeT5A (Supplementary Fig. 1f). These data together support the specific involvement of P-cadherin during HM-MeT5A interactions.

We next investigated the mechanical properties of P-cadherin during the adhesion using atomic force microscopy-based single-cell spectroscopy to measure the force of interaction between MeT5A cells cultured on plates and ovarian cancer cells attached to the cantilever (Fig. 1d). HM cells exhibited a significantly higher rupture force compared to NM cells (Fig. 1e), indicating the presence of stronger mechanical stress at the junctions between HM and MeT5A cells in contrast to the junctions between NM and MeT5A cells. The rupture force in HM–MeT5A interactions decreased when treated with siP-cad or a P-cadherin neutralizing antibody, compared to interactions treated with siNS or control IgG treatment, respectively (Fig. 1f). These findings suggest that P-cadherin plays an essential role in mediating mechano-interactions between these two cell types. Notably, we also observed enhanced P-cadherin expression in ascitic tumor cells derived from ovarian cancer patients compared to normal ovarian surface epithelium (OSE) (Fig. 1g). To test the functional role of P-cadherin, we further used magnetic beads coated with immobilized P-cadherin Fc chimera protein (P-cad/Fc), which bind to the cell surface and mimic cadherin-mediated adhesion^23^ (Fig. 1h). The presence of P-cad/Fc-coated beads significantly enhanced cell proliferation in HM cells, whereas IgG Fc-coated beads showed no effect when compared with the untreated control (Fig. 1i). These findings suggest a potential role of P-cadherin in promoting tumor growth.

### P-cadherin rewires lipid metabolism in tumor cells

Metabolic reprogramming enables cancer cells to meet the increased demand for energy and biosynthetic precursors necessary for their uncontrolled growth during metastasis. The observed enhancement in proliferation resulting from P-cadherin-coated beads prompted us to hypothesize that P-cadherin might affect cellular metabolism. To test this, we performed RNA-seq analysis of HM cells upon treatment with siNS or siP-cad. Pathway analysis revealed a significant alteration in fatty acid (FA) metabolism (*P* < 0.05, Fig. 2a). To validate our RNA-seq findings, we performed RT-PCR for several key lipogenic genes, namely acetyl-CoA acetyltransferase 2 (ACAT2), ATP-citrate lyase (ACLY), FA synthase (FASN), lipin 1 (LPIN1), and sterol regulatory element-binding transcription factor 1 (SREBF1). All of these genes exhibited significant decreased expression levels in HM cells treated with siP-cad (Fig. 2b). Consistently, the overexpression of P-cadherin in NM cells led to the upregulated expression of these lipogenic genes (Fig. 2b). Western blot analysis further confirmed similar changes at the protein level (Supplementary Fig. 2a). We also tested the expression of well-known lipolytic genes, including hormone-sensitive lipase (HSL), monoacylglycerol lipase (MAGL) and perilipin-1 (PLIN), which remained unchanged regardless of P-cadherin expression (Fig. 2b, Supplementary Fig. 2a), confirming that P-cadherin specifically regulates lipogenesis rather than lipolysis in HM cells. Moreover, HM cells sorted from the MeT5A coculture showed significant upregulation of the above key lipogenic proteins compared to HM cells cultured alone (Fig. 2c and Supplementary Fig. 2b). This was supported by Oil Red O (ORO) staining, which revealed greater accumulation of neutral lipids in the HM-MeT5A coculture than in HM cells alone, whereas no change was observed in NM cells upon coculture (Fig. 2d and Supplementary Fig. 2c). Compared with IgG and siNS controls, P-cadherin antibody and siP-cad markedly impaired the lipid accumulation in HM cells, as revealed by decreased ORO staining in the HM–MeT5A coculture (Fig. 2e and Supplementary Fig. 2d). Consistently, overexpression of P-cadherin in NM enhanced lipid accumulation upon coculture (Fig. 2f). These observations collectively suggest a specific role of P-cadherin in regulating *de novo* lipogenesis in HM cells during mesothelial adhesion. We also pretreated HM cells with 5-(tetradecyloxy)-2-furoic acid, which inhibits acetyl-CoA carboxylase essential for FA synthesis^23^ or silenced SREBP1, which is a master regulator of lipogenesis^24^. Both treatments resulted in a significant decrease in cellular lipid content during the HM–MeT5A coculture (Fig. 2g and 2h), suggesting that additional lipids are produced through *de novo* lipogenesis instead of being transferred from the extracellular environment.

**Fig. 2.**
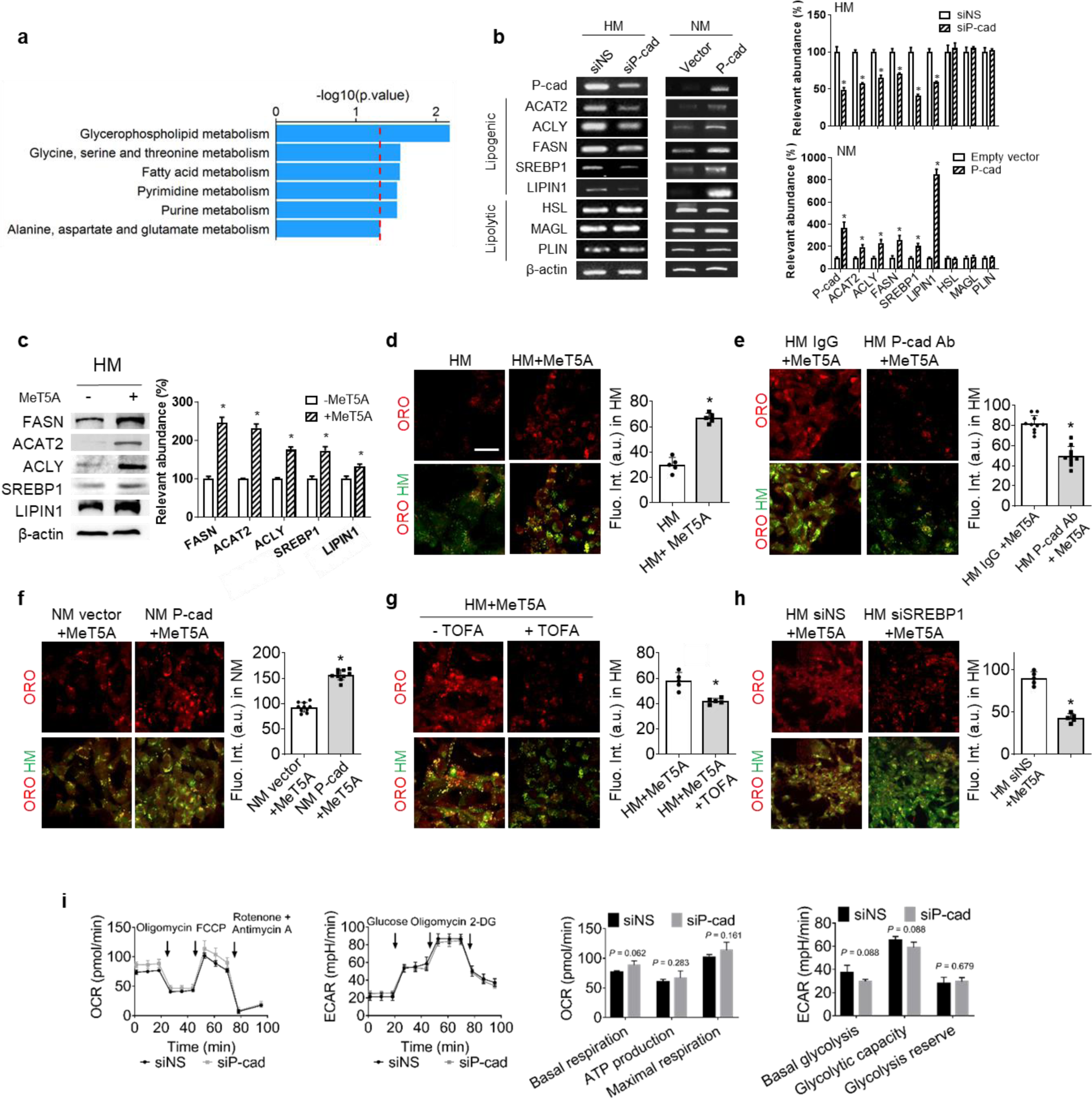
P-cadherin rewires lipid metabolism in tumor cells. **a,** RNA-sequencing analysis was performed with siNS or siP-cad-treated HM cells. Pathways downregulated in siP-cad compared to siNS are shown. Red line means P value = 0.05. **b,** RT-PCR analysis of the expression of P-cadherin (P-cad), lipogenic genes (ACAT2, ACLY, FASN, SREBP1 and LIPIN1) and lipolytic genes (HSL, MAGL and PLIN) in siNS, siP-cad, empty vector or P-cad overexpression vector-transfected HM cells. **c,** Western blot of FASN, ACAT2, ACLY, SREBP1 and LIPIN1 in HM cells cultured alone or cocultured with MeT5A. b-c, β-actin serves as the loading control. Data are presented as mean ± SD (n=3). **d-h,** Representative images and quantification of oil red O (ORO) staining of neutral lipids (red) in CMFDA-labelled HM or NM cells (green) cocultured with MeT5A for 24 hours. Scale bar, 20 μm. At least five microscopic fields were quantified for each condition and data are presented as mean ± SD. **d,** HM cells were cultured alone or cocultured with MeT5A for 24 hours. **e,** HM cells treated with P-cadherin antibody (P-cad Ab) or IgG isotype control before MeT5A coculture. **f,** NM cells transfected with P-cad overexpression vector or empty vector control before MeT5A coculture. g, HM cells were pretreated with or without the acetyl-CoA carboxylase-α inhibitor TOFA (20 μM) before MeT5A coculture. **h,** HM cells were transfected with siNS or SREBP1 siRNA (siSREBP1) before MeT5A coculture. **i**, Mitochondrial metabolism (left panel) and glycolysis profiles (right panel) of HM cells treated with siNS or siP-cad and corresponding quantification of bioenergetic masses (n= 4 per group). OCR: Oxygen consumption rate; ECAR: extracellular acidic rate. Acquisition settings remained the same in all groups of fluorescent images. Data are presented as mean ± SD. *P < 0.05.

We also performed Seahorse XF analysis to assess the influence of P-cadherin on bioenergetic metabolism in HM cells. Our results showed that siP-cad did not alter mitochondrial or glycolytic metabolism in HM cells (Fig. 2i). Consistently, RT-PCR showed that siP-cad had no effect on the uptake of glucose or the expression of glycolytic genes, including glucose transporter 1 (GLUT1), hexokinase 2 (HK2), glucose-6-phosphate isomerase (GPI), and phosphoglycerate kinase 1 (PGK1), in HM cells (Supplementary Fig. 2e and 2f). Gene set enrichment analysis (GSEA) of RNA-seq data also suggested that the lipolytic or glycolytic pathways were unaffected (Supplementary Fig. 2g). Together, these results further indicate that P-cadherin-mediated tumor– mesothelium adhesion specifically evokes *de novo* lipogenesis in HM cells.

### P-cadherin-mediated mechanotransduction regulates lipogenesis through SREBP1 activation

We next investigated the mechanistic basis for the link between P-cadherin-mediated mechanotransduction and cell metabolism. Among the lipogenic genes upregulated by P-cadherin, we chose to focus on SREBP1 (encoded by the *SREBF1* gene) since it is an important transcription factor that regulates FA synthesis and has been shown to be mechanically regulated by extracellular matrix stiffness^24,25^. Treatment with P-cad/Fc beads led to the upregulation of SREBP1 mRNA as well as both the full-length (fl) precursor and mature (ma) forms of SREBP1 proteins in HM cells (Fig. 3a). Immunostaining showed enhanced nuclear levels of SREBP1 upon P-cad/Fc bead treatment, an effect that could be inhibited by the P-cadherin blocking antibody (Fig. 3b), confirming the specific role of P-cadherin in SREBP1 activation. The expression of lipogenic genes induced by P-cad/Fc-coated beads was found to be suppressed upon knockdown of SREBP1 (Fig. 3c). Furthermore, the reduction in lipogenic gene expression caused by siP-cad could be restored by overexpressing SREBP1 (Fig. 3d), providing further confirmation of the involvement of SREBP1 in P-cadherin-mediated lipogenesis. To investigate how P-cadherin transduces this signal, we evaluated the effects of mechanical cues on the actin cytoskeleton. Phalloidin staining showed that P-cadherin-mediated adhesion force significantly enhanced filamentous actin tension, which was blocked by treatment with P-cadherin antibody (Fig. 3e and 3f). Treatment of HM cells with blebbistatin, an inhibitor of myosin that disrupts actin organization, inhibited the induction of SREBP1 by P-cad/Fc-coated beads (Fig. 3g).

**Fig. 3.**
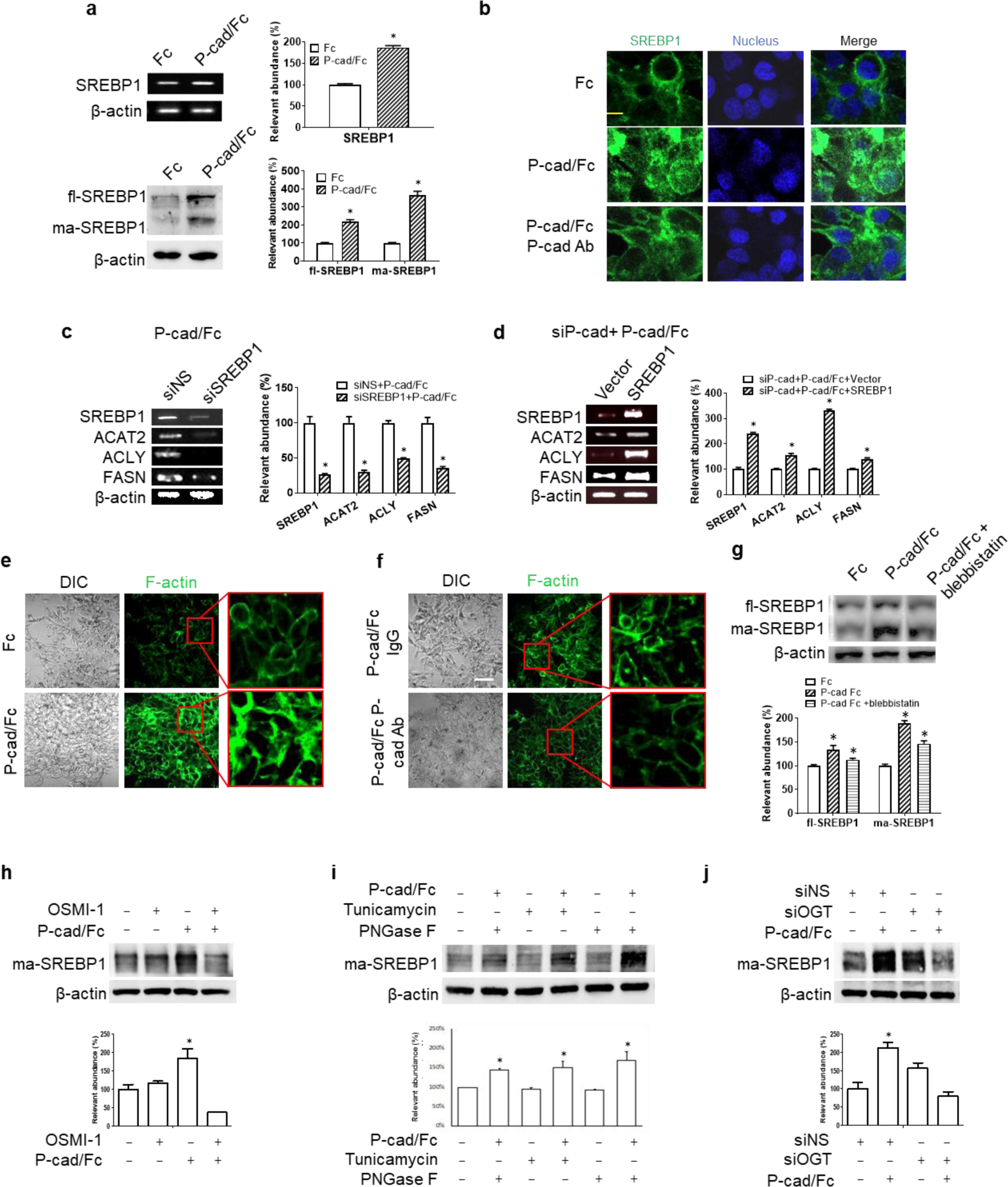
P-cadherin mechano-activates SREBP1 via actin remodeling in a O-GlcNacylation-dependent manner. **a,** HM cells were treated with IgG/Fc (Fc)- or P-cad/Fc-coated beads for 24 h, followed by RT-PCR or Western blot analysis of SREBP1. **b,** HM cells were incubated with Fc- or P-cad/Fc-coated beads in the absence or presence of P-cadherin blocking antibody (P-cad Ab) for 30 min, followed by immunostaining of SREBP1. Scale bar, 20 μm. **c,** RT-PCR analysis of SREBP1, ACAT2, ACLY and FASN in HM cells transfected with siNS or siSREBP1 in the presence of P-cad/Fc-coated beads. **d,** HM cells were coo-transfected with siP-cad and empty vector or SREBP1 overexpression vector, followed by incubation with P-cad/Fc-coated beads. RT-PCR analysis of the indicated genes was performed. **e-f,** Phalloidin staining of F-actin in HM cells treated with Fc or P-cad/Fc coated beads **(e)** and in HM cells treated with or without P-cad antibody in the presence of P-cad/Fc-coated beads **(f)**. **g,** Expression of full-length (fl)- and mature(ma)-SREBP1 in HM cells with the indicated treatment were analyzed by Western blot analysis. **h,** HM cells were treated with beads coated with Fc or P-cad/Fc and OSMI1 (10 μM) as indicated, followed by Western blot analysis. **i,** HM were treated with tunicamycin (100 nM), PNGase F and Fc or P-cad/Fc-coated beads and as indicated, followed by Western blot analysis. **j,** HM cells transfected with siNS or OGT siRNA (siOGT) were treated with Fc or P-cad/Fc-coated beads, followed by Western blot analysis. β-actin serves as the loading control for both RT-PCR and Western blot. Data are presented as mean ± SD from 3 independent experiments. \**P* < 0.05.

Cell adhesion is controlled by dynamic posttranslational modifications (PTMs) that facilitate rapid biological responses to stimuli. O-GlcNAcylation, a widespread PTM that acts as an important sensor of metabolic state, has recently been linked to lipogenesis and SREBP1 regulation^27,28^. Therefore, we asked if it is involved in P-cadherin signaling. Treating HM cells with OSMI-1, an inhibitor of O-GlcNAc transferase (OGT), suppressed SREBP1 induction by P-cad/Fc coated beads (Fig. 3h). However, both treatment with tunicamycin, an inhibitor of N-glycan biosynthesis, and surface N-glycan removal by PNGase F had no effect on the induction of SREBP1 by P-cad/Fc-coated beads (Fig. 3i). Consistent with OSMI-1 treatment, knockdown of OGT by an siRNA inhibited SREBP1 induction (Fig. 3j). These data suggest that P-cadherin regulates SREBP1 in an O-GlcNAcylation-dependent manner.

### P-cadherin activates glycolysis in mesothelial cells

We next determined whether tumor-mesothelial interaction could induce changes in the activity of mesothelial cells. Compared with MeT5A alone, HM, but not NM, cells selectively activated MeT5A to secrete tumor-promoting cytokines, such as G-CSF, GM-CSF, IL-6, IL-8, CXCL1/GROα, and IL-1α/IL-1F1 (Fig. 4a). As shown in Fig. 4b, we noted a marked decrease in E-cadherin expression and a strong induction of N-cadherin and fibronectin expression in MeT5A sorted from the HM coculture compared with MeT5A alone, suggesting the presence of mesothelial-to-mesenchymal transition that is indicative of mesothelial cell activation^6^. Similar results were observed using HPMC (Supplementary Fig. 3a).

**Fig. 4.**
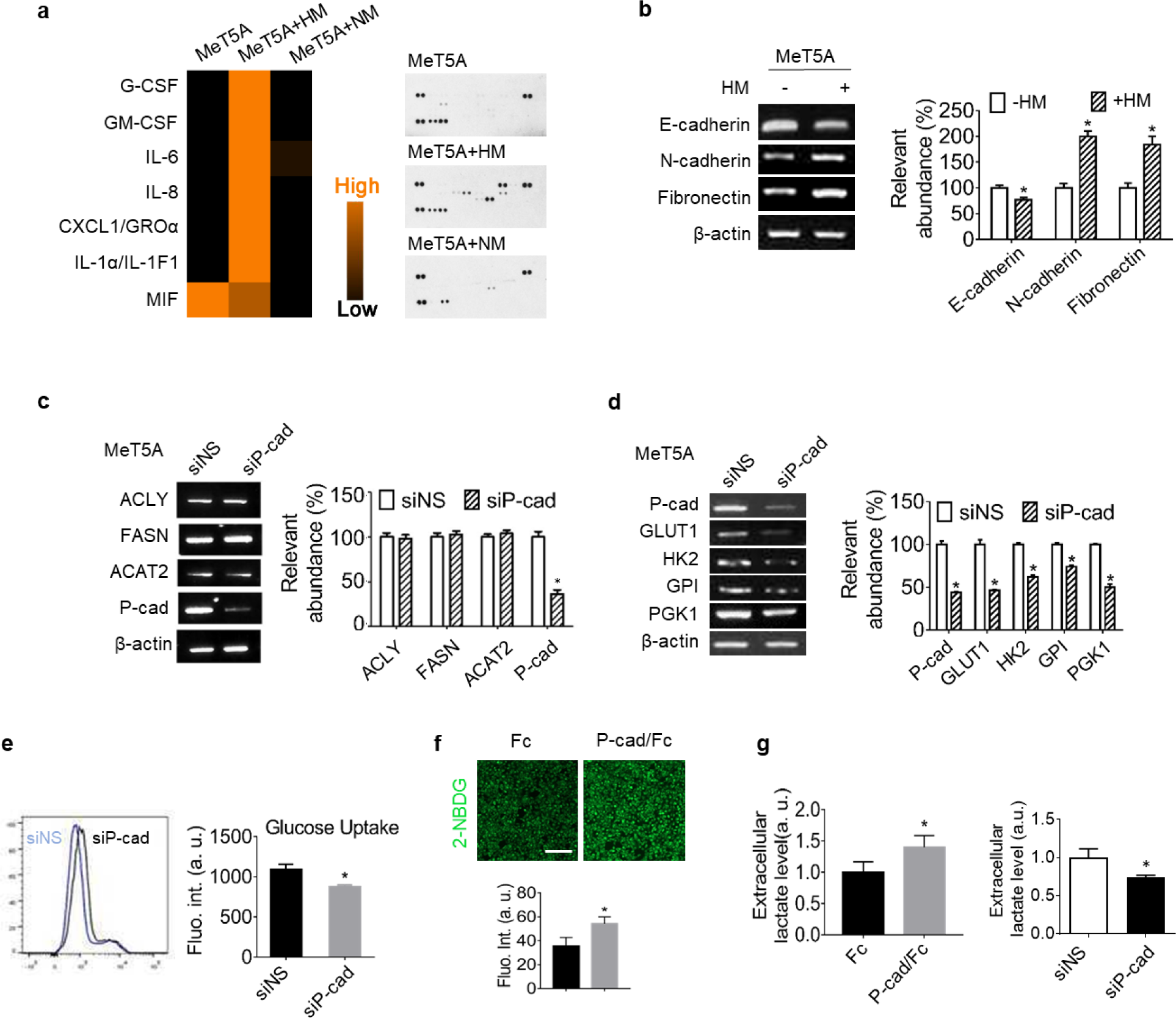
P-cadherin regulates glycolysis in MeT5A cells. **a,** Cytokine array analysis of conditioned medium from MeT5A cells alone or cocultured with HM or NM cells. **b,** RT-PCR analysis of E-cadherin, N-cadherin, and fibronectin levels in MeT5A cocultured with or without HM cells were shown. **c-d,** MeT5A were transfected with siNS or siP-cad and cocultured with HM. After coculture, MeT5A were sorted by flow cytometry followed by RT-PCR for **(c)** P-cad and lipogenic genes (ACAT2, ACLY and FASN) and **(d)** P-cad and glycolytic genes (GLUT1, HK2, GPI and PGK1). β-actin serves as the loading control. **e,** Glucose uptake in siNS or siP-cad-treated MeT5A cells was analyzed by flow cytometry using 2-NBDG, a fluorescent D-glucose analog. **f,** Representative fluorescent images of 2-NBDG uptake by MeT5A cells treated with Fc or P-cad/Fc coated beads for 24 h were shown, and fluorescent intensity were quantified (n= 10). **g,** Extracellular lactate levels in the conditioned medium of MeT5A cells treated with Fc- or P-cad/Fc-coated beads for 24 h (left) and siNS or siP-cad (right) was measured. Data are presented as mean ± SD from 3 independent experiments. \**P* < 0.05.

Considering the impact of P-cadherin on HM metabolism, it is plausible that P-cadherin might also influence the metabolic profile of mesothelial cells. However, upon silencing P-cadherin, we observed no effect on the expression levels of lipogenic genes (Fig. 4c and Supplementary Fig. 3b). Interestingly, we discovered that treatment with siP-cad or P-cadherin overexpression led to a significant decrease or increase, respectively, in key glycolytic genes (GLUT1, HK2, GPI, and PGK1) in mesothelial cells (Fig. 4d and Supplementary Fig. 3c, 3d). Corresponding changes at the protein level were observed by Western blot analysis (Supplementary Fig. 3b-d). Additionally, flow cytometric analysis revealed a reduced capacity of MeT5A cells to uptake glucose, as evidenced by a decrease in the fluorescence signal of 2-NBDG (a glucose analogue) following siP-cad transfection compared to siNS transfection (Fig. 4e). In contrast, treatment of MeT5A cells with P-cad/Fc-coated beads led to a significant increase in glucose uptake and subsequent lactate production, which was reversed by siP-cad treatment (Fig. 4f and 4g). These findings provide compelling evidence that P-cadherin-mediated adhesion forces promote a highly glycolytic profile in mesothelial cells.

### Lactate is shuttled to HM cells as a lipid source

Given the differential metabolic alterations mediated by P-cadherin in tumor and mesothelial cells, we postulated that the lactate produced by the activated mesothelium may drive HM tumor cell lipogenesis. We first traced the lipidomic fate of ^13^C-labeled lactate in metastatic tumors in mice through GC-MS/MS and found that palmitic acid (C16:0) in HM tumor cells was labeled to a larger degree (M+2 to M+8) than that in NM tumor cells (M+2 to M+4), indicating that HM cells use more lactate for *de novo* FA synthesis compared to NM cells (Supplementary Fig. 4a). To provide direct evidence that mesothelial cell-derived lactate could promote HM lipogenesis, we performed targeted lipidomics in the HM-MeT5A coculture after incubation with uniformly ^13^C-labeled glucose ([U^13^C_6_]-glucose) with or without the addition of cold lactate (Fig. 5a). In the presence of cold lactate, the generation of high ^13^C-labeled palmitic acids was shifted to low ^13^C-labeled palmitic acid (16:0) in the sorted HM cells (Fig. 5b). Similar results were observed for palmitoleic acid (16:1) and cholesterol synthesis (Fig. 5b), confirming that MeT5A-secreted lactate is used by HM cells for lipid generation (Fig. 5c). Consistently, siP-cad reduced the lactate uptake of HM induced by P-cad/Fc-coated beads (Supplementary Fig. 4b).

**Fig. 5.**
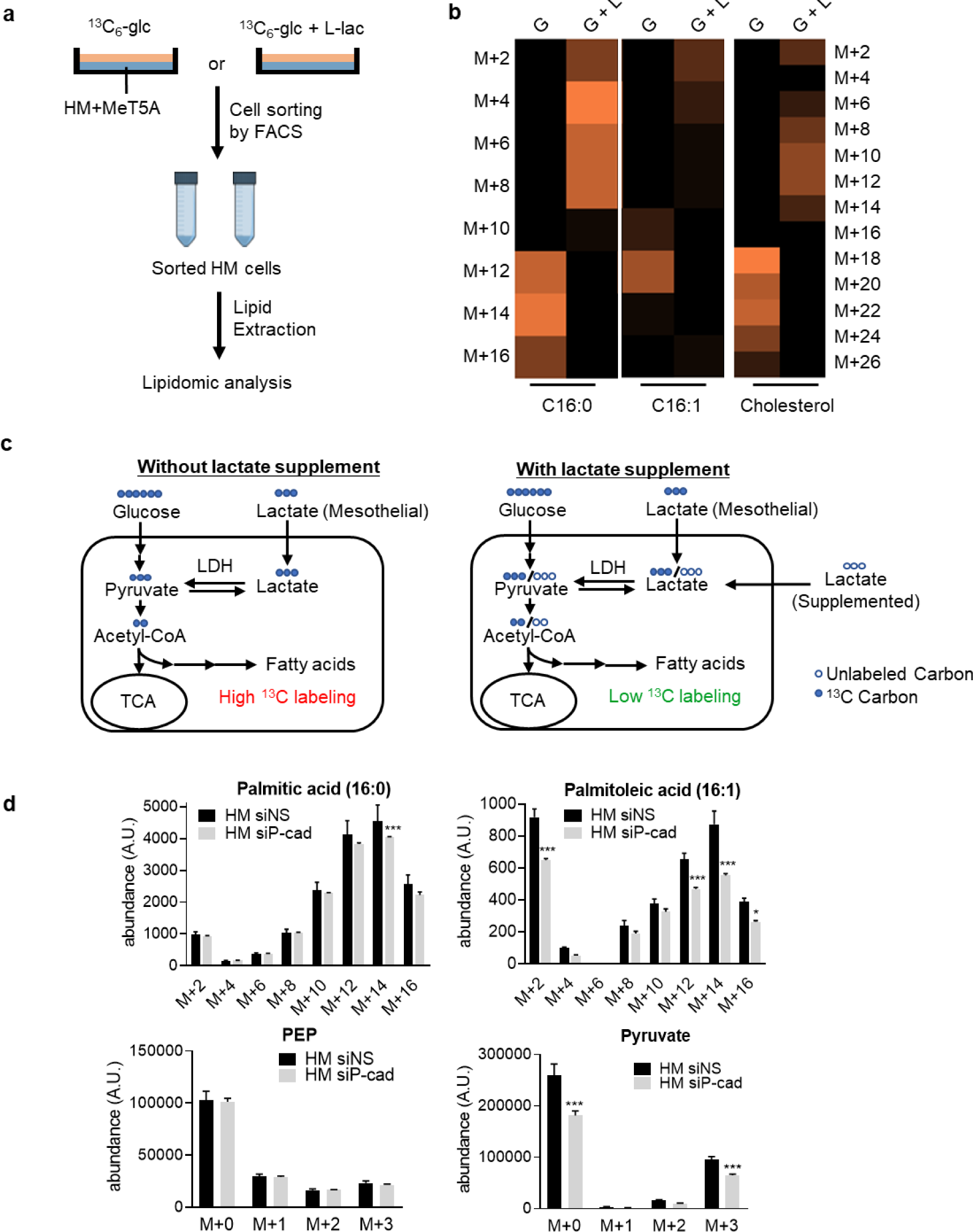
Lactate is shuttled from mesothelial cells into HM cells. HM and MeT5A were cocultured in medium supplemented with 25 mM [U^13^C]-glucose (G) or 25 mM [U^13^C]-glucose and 11 mM L-lactate (G+L) for 24 h, followed by targeted lipidomic analysis. **a,** Schematic illustration of the ^13^C-labeled targeted lipidomic assay workflow. **b,** Heatmap of ^13^C-labeled fatty acids and cholesterol (C16:0, C16:1, CHOL) in sorted HM was shown. **c,** Schematic illustration of the expected effects of supplemented cold lactate acid on lipid composition. **d,** Lipidomic analysis of ^13^C-labeled palmitic acid (C16:0), palmitoleic acid (C16:1), glycolysis intermediates PEP and pyruvate in HM cells with siNS or siP-cad after coculture with MeT5A in medium supplemented with 25 mM [U^13^C]-glucose (with three technical replicates each). Data are presented as mean ± SD. \**P* < 0.05, \*\*\**P* < 0.001 Two way ANOVA test. PEP, phosphoenolpyruvate.

We also performed ^13^C glucose experiment to confirm the role of P-cadherin in lactate shuttling during coculture. Comparing to siNS control, siP-cad caused a significant decrease in ^13^C-labelled palmitic acid (M+14) and palmitoleic acid (M+12 to M+16) in HM cells, suggesting a role of P-cadherin in lipogenesis (Fig. 5d). There was also a significant reduction in ^13^C-labelled pyruvate, while no difference was found in phosphoenolpyruvate, a metabolic product upstream of pyruvate in the glycolytic pathway (Fig. 5d). These data support our results that P-cadherin did not affect the glycolysis pathway but may affect the use of ^13^C-labeled lactate from MeT5A as a source for lipogenesis.

We hypothesized that lactate shuttling is important for tumor–mesothelial interaction. Monocarboxylate transporter 4 (MCT4) and monocarboxylate transporter 1 (MCT1) are the major transporters of lactate efflux and uptake, respectively^29,30^. As shown in Fig. 6a and 6b, suppressing lactate export in MeT5A with siMCT4 restricted both lipid generation and HM cell proliferation during coculture. Alternatively, inhibiting lactate import by MCT1 shRNA in HM cells abolished lipid accumulation and proliferation (Fig. 6c and 6d). AR-C155858^31^, a MCT1/2 inhibitor, was also used to block lactate uptake in HM cells. AR-C155858 significantly reduced the ORO staining in the HM–MeT5A coculture, further confirming that lactate shuttling is required for lipogenesis in tumor cells (Fig. 6e). These results demonstrate that P-cadherin-mediated adhesion drives *de novo* lipogenesis in highly metastatic tumor cells through the vectorial transport of lactate from the mesothelium to cancer cell.

**Fig. 6.**
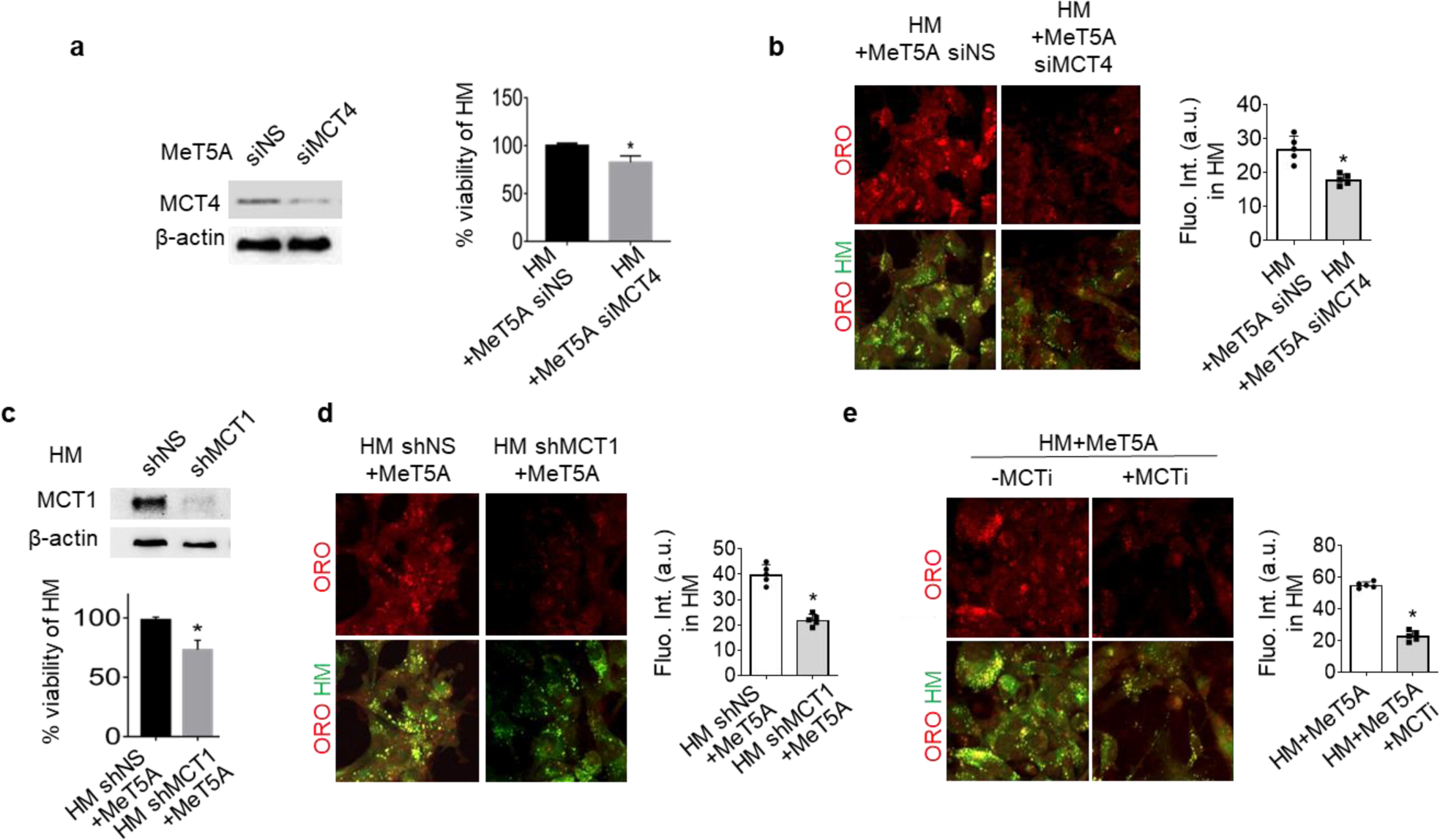
Lactate shuttling via MCT4 in mesothelial cells and MCT1 in HM cells promotes lipogenesis. **a,** Western blot analysis of MCT4 in MeT5A cells transfected with siNS or MCT4 siRNA (siMCT4). **b**, Representative images and quantification of oil red O (ORO) staining in CMFDA-labelled HM (green) cocultured with MeT5A transfected with siNS or siMCT4 were shown. **c**, Western blot analysis of MCT1 in HM cells stably transfected with non-specific or MCT1 shRNA (shNS or shMCT1). Viability of HM cells with shNS or shMCT in MeT5A coculture was shown. **d**, HM with non-specific or MCT1 shRNA (shNS or shMCT1) were cocultured with MeT5A. Representative images and quantification of ORO staining (red) in CMFDA-labelled HM (green) were shown. *P < 0.05. **e**, Representative images and quantification of ORO staining (red) in CMFDA-labelled HM (green) in the absence or presence of monocarboxylate transporter inhibitor (MCTi) for 24 h and MeT5A coculture were shown. Scale bar, 20 μm. β-actin was used as the loading control for all Western blots (n=3). Data are presented as mean±SD. *P < 0.05.

### High FASN expression is associated with high P-cadherin and Ki67 expression in metastatic ovarian cancer

To corroborate the clinical relevance of lipogenesis in peritoneal metastasis, we first performed immunohistochemical staining of FASN in paired primary and metastatic tumor samples obtained from HGSOC patients. The metastatic samples include omental metastases as well as metastatic specimens from other sites with mesothelial lining (Total n=16: omentum, n=9; bladder flap, n=4; pelvic peritoneum/peritoneal, n=3). The clinicopathological information is provided in Supplementary Table 1. We specifically examined the FASN expression in tumor cells rather than adipocytes, which are histologically distinct. As shown in Fig. 7a and 7b, the majority of metastatic specimens expressed elevated levels of tumor FASN (H-score) as compared to their parallel primary samples (10 of 16 metastatic specimens, 62.5%, *P*=0.0362). To validate the association between FASN, P-cadherin and metastatic growth, we further evaluated the expression levels of P-cadherin and the proliferation marker Ki67. Metastatic specimens with elevated FASN expression showed higher expression of P-cadherin and Ki67 compared to matched primary tumors (P-cadherin, *P*=0.0340; Ki67, *P*=0.0241), while metastatic specimens with low FASN expression showed little changes in P-cadherin and Ki67 expression (Fig. 7c and 7d). The higher expression levels of FASN, P-cadherin and Ki67 in HM relative to NM tumors were also confirmed (Supplementary Fig. 5a and 5b). These data support the association of P-cadherin with cell proliferation through lipogenic regulation.

**Fig. 7.**
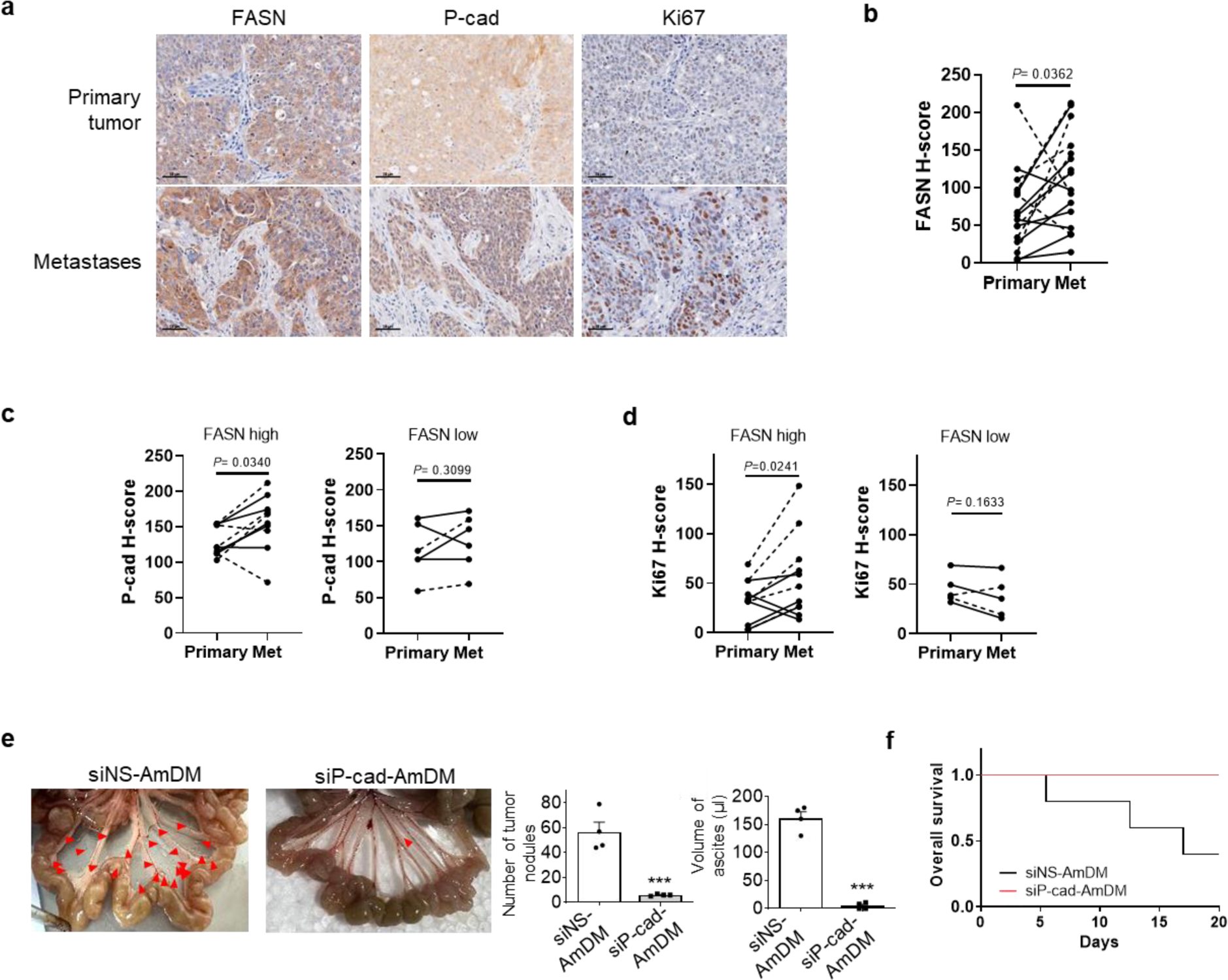
High FASN expression is associated with high P-cadherin and Ki67 expression in metastatic ovarian cancer. Immunohistochemical (IHC) analysis of FASN, P-cadherin (P-cad) and the proliferation marker Ki67 expression in primary patient specimens (n= 10) and paired metastatic specimens of omental and other peritoneal sites (n= 16). **a,** Representative images of IHC staining of FASN, P-cad and Ki67. **b,** Quantitative analysis (H-score) of FASN expression in paired primary and metastatic specimens of omental (solid lines) and other peritoneal sites (dashed lines). **c-d,** Quantitative analysis of P-cad **(c)** or Ki67 **(d)** expression in patients with elevated metastatic FASN expression vs low metastatic FASN expression. Data presented are median value. Paired t-test, *P*< 0.05 is considered significant. **e-f,** Mice were intraperitoneally injected with HM cells followed by intravenous injection of siNS-AmDM or siP-cad-AmDM for 3 weeks (siRNA: 1.0 mg/kg, N/P ratio=5, twice a week). Representative images of the peritoneal metastases were shown. Tumor nodules and ascites volume were quantified (n= 5) **(e)**. Data are presented as mean ± SD. \*\*\**P* < 0.001. **f,** Kaplen-Meier curves show survival of mice with siNS-AmDM and siP-cad-AmDM treatment.

To confirm the importance of P-cadherin signaling *in vivo*, we used our well-characterized amphiphilic dendrimers, AmDM, which formed nanoparticles with siRNA and could specifically target tumor sites through enhanced permeation and retention effects^32,33^. Mice with intraperitoneal xenograft of HM were treated with dendriplexes of siP-cad, which caused significant reductions in the number of metastatic tumor nodules and ascites formation in the peritoneal cavity, as well as enhanced survival as compared with mice injected with control dendriplexes of siNS (Fig. 7e and 7f). A representative high magnification image of the tumor nodules and histological H&E staining were shown in Supplementary Fig. 6a and 6b to confirm the identification of metastatic tumors. Successful P-cadherin knockdown using dendriplexes was confirmed by Western blot (Supplementary Fig. 6c). These data validated an important role of P-cadherin during metastasis.

### Inhibition of MCT1/4 reduces peritoneal metastasis *in vivo*

High expression of MCT1 was found to be significantly associated with a decrease in both overall survival and progression-free survival of ovarian cancer patients using Kaplan-Meier Plotter (*P* < 0.005) (Fig. 8a). MCT1 also showed a positive correlation with P-cadherin and FASN expression (Fig. 8b), implicating the importance of the lactate uptake. To further confirm lactate shuttling *in vivo*, mice bearing intraperitoneal HM tumors were treated with dendriplexes of siMCT1 or siMCT4. Successful target gene knockdown using dendriplexes was confirmed by Western blot analysis (Supplementary Fig. 6c). We observed significant reductions in the number of metastatic tumor nodules and ascites formation compared to mice injected with siNS-dendriplexes control (Fig. 8c and 8e). Kaplan-Meier curves showed that mice with specific targeting of MCT1 or MCT4 also displayed prolonged survival (Fig. 8d and 8f). The importance of MCT1 in peritoneal metastasis was also supported by shRNA knockdown (Supplementary Fig. 6d). We further demonstrated the potential of therapeutic targeting by treatment with a clinical grade MCT1 inhibitor, AZD3965^34^, which significantly reduced metastasis in mice (Supplementary Fig. 6e). These *in vivo* experiments, together with patient analysis, strongly support the clinical importance of our proposed mechanisms in ovarian cancer metastasis.

**Fig. 8.**
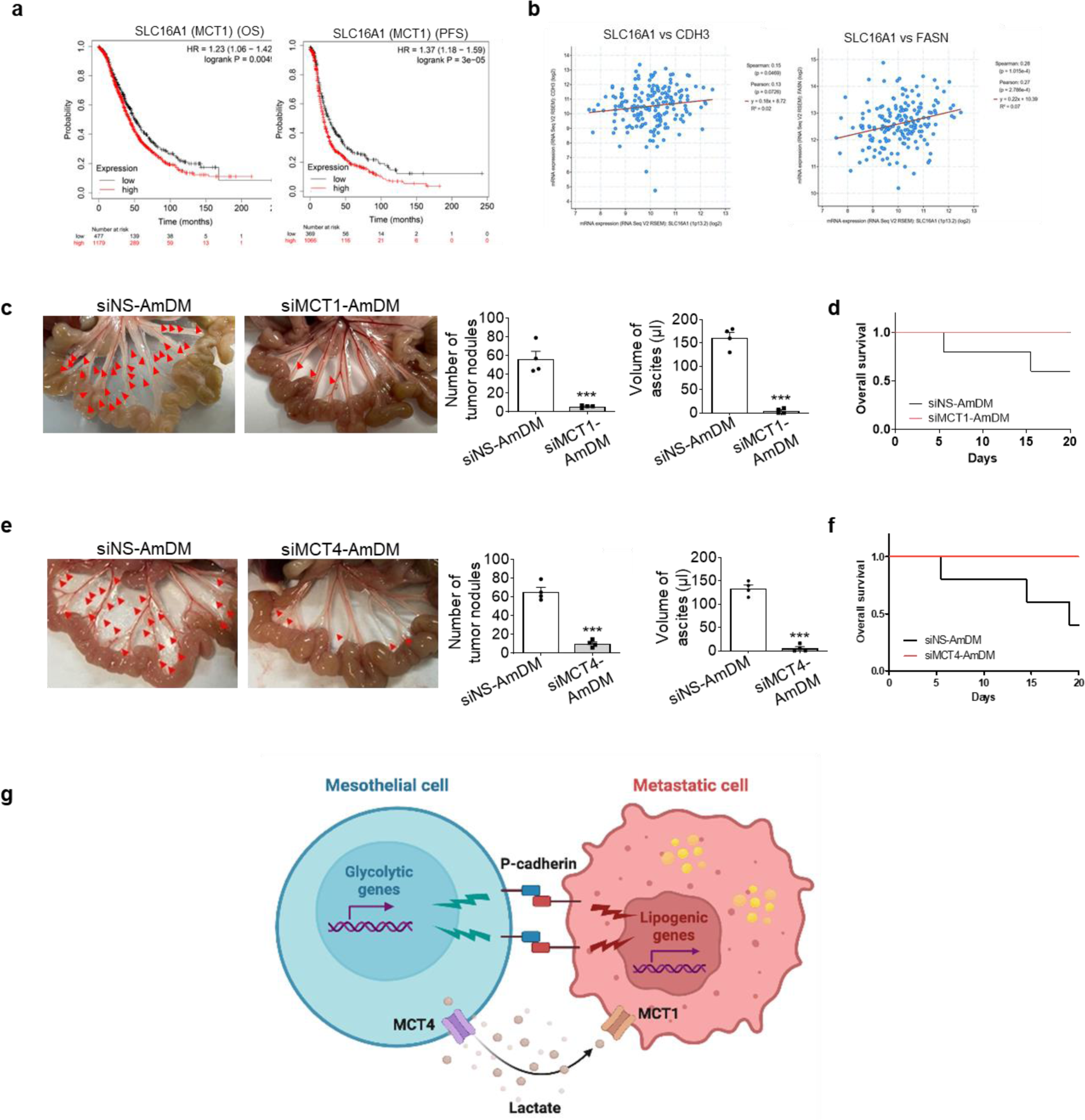
MCT1/MCT4-mediated lactate shuttling is a potential therapeutic target for peritoneal metastasis. **a,** Kaplan-Meier plot of overall survival (OS) and progression-free survival (PFS) in ovarian cancer patients with low and high expression of MCT1. Number at risk is listed below. **b,** SLC16A1 (MCT1) mRNA transcript level positively correlates with CDH3 (P-cadherin) and FASN transcript levels. **c-f,** Mice bearing intraperitoneally injected HM cells were intravenously injected with siNS-AmDM or siMCT1-AmDM **(c,d)** and siNS-AmDM or siMCT4-AmDM **(e,f)** for 3 weeks (siRNA: 1.0 mg/kg, N/P ratio=5, twice a week). Representative images of the peritoneal metastases were shown. The number of metastatic nodules and ascites volume was quantified (n= 4). **d,f,** Kaplen-Meier curves show overall survival of mice with siNS-AmDM, siMCT1-AmDM or siMCT4-AmDM treatment as indicated. **g,** Schematic illustration of the proposed mesothelial-metastatic cell metabolic coupling through P-cadherin. Data are expressed as mean ± SEM. ***, *P* <0.001.

## Discussion

Adhesion to the mesothelium and the ability to sustain proliferation at the secondary site are pivotal events in peritoneal metastasis; however, the underlying mechanisms remain poorly elucidated. In this study, we have identified P-cadherin as a distinct and critical mediator that initiates metabolic symbiosis between tumor cells and the interacting mesothelium (Fig. 8g). These findings are consistent with preclinical and clinical evidence which suggest a correlation between high expression of P-cadherin, metastasis, and unfavorable clinical outcomes in ovarian cancer^35,36,37,38^, thereby emphasizing the therapeutic potential of targeting P-cadherin in the management of peritoneal metastasis. While cadherin-mediated adhesion typically occurs within a complex milieu where diverse cell types engage in physical interactions, the majority of cadherin studies have predominantly focused on homotypic interactions among cells of the same lineage. In contrast, our results clearly show that homophilic interactions mediated by P-cadherin could elicit distinct signaling responses in tumor and mesothelial cells at the secondary site. This novel finding provides invaluable insights into the spatial dynamics of cadherin adhesion and signaling during the metastatic process. Moreover, this specific metabolic coupling may underlie the organotropism of metastatic cells towards the peritoneum, potentially influencing the prognosis and treatment strategies for cancer patients.

How mechanical pathways are integrated with other cellular processes, such as cell metabolism, remains largely unknown. P-cadherin was shown to be associated with a hypoxic and glycolytic phenotype characterized by the expression of HIF-1α, GLUT1, CAIX, MCT1, and CD147^39^. In addition, P-cadherin promoted anoikis resistance by activating the pentose–phosphate pathway and reducing oxidative stress in breast cancer^40^. However, the precise molecular mechanisms underlying these processes remain unidentified. Our study revealed that P-cadherin induces lipogenesis in HM cells by facilitating actin remodeling and activating SREBP1. Interestingly, previous research has shown that elevated extracellular matrix stiffness inhibits SREBP1 activity through increased acto-myosin contraction and AMPK activation^25^. This discrepancy in signaling could be attributed to the distinct mechanotransduction processes occurring at context-dependent cell-matrix and cell-cell adhesion sites^41^. Nonetheless, high expression of SREBP1 has been associated with advanced disease stages and poor prognosis in ovarian cancer and other cancer types^24,42^, which is in line with our observations. Conversely, we found that HM cells could enhance glycolysis in the neighboring mesothelium. In addition to serving as a source of lactate, the hyperglycolytic phenotype of the mesothelium has been reported to promote mesothelial-to-mesenchymal transition and peritoneal fibrogenesis^43^, thereby reinforcing the metastatic process.

Lipids play a causal role in driving tumorigenesis and metastasis^44^. Overexpression of lipogenic genes has been associated with metastatic progression and poor prognosis in many cancer types. In ovarian cancer, the increased expression of FASN is observed in tumor tissues and predicts advanced disease status and poor clinical outcomes^45^. Inhibition of lipid metabolism using an inhibitor cocktail was shown to reduce the peritoneal metastasis of ovarian cancer *in vivo*^46^. Consistent with these findings, our data suggest that a P-cadherin-mediated lipogenic phenotype characterized by the overexpression of lipogenic genes is important for metastasis *in vivo*. Beyond serving as an energy source or building blocks for membrane synthesis, lipids play other important roles in the tumor microenvironment. For example, FAs and cholesterol enhance therapeutic resistance and impact immune cell functions^47,48^. The lipid pool in cancer cells can be supplied by both intracellular and extracellular sources, although the exact stoichiometric relationships between these sources remain to be determined^49^. Various cancers, including ovarian cancer, have been shown to uptake FAs produced by surrounding adipose tissues^50^. However, in some contexts, exogenously supplemented FAs could not compensate for *de novo* synthesized FAs, such as through autophagy in adipocytes and mitosis in cancer cells^51,52^, suggesting that *de novo* lipogenesis is required for unique cellular processes. Furthermore, *de novo* lipogenesis may enable cancer cells to tailor their FA composition to meet their unique requirements. Notable differences in FA composition have been observed between omental fat deposits and malignant ovarian tumors. In addition, FA species vary in their length and saturation, and such diversity indicates that their availability depends on the specificity of metabolic pathways. Recent studies have shown that unsaturated FAs were enriched in ovarian cancer stem cells and were essential for their proliferation and survival^53^. Therefore, *de novo* lipogenesis may enable cancer cells to tightly control their FA composition. Moreover, metabolic intermediates in *de novo* lipogenesis (citrate, acetyl-CoA, malonyl-CoA, and palmitate) can function as secondary messengers for other signaling pathways^54^. Thus, the high lipid demand for metastatic growth is not only met by the tumor microenvironment but also by cancer cells themselves, making lipogenesis an attractive target for therapy.

Emerging evidence suggests lactate as an oncometabolite in the tumor microenvironment. For example, lactate was shown to promote the immunosuppressive function of Treg cells through lactylation^55^. Significantly higher lactate levels were found in ovarian cancer peritoneal metastasis samples than in benign samples in a small cohort of patients^56^. Furthermore, high lactate transporter expression predicted advanced disease status and poor clinical outcomes in ovarian cancer^57^. Our study has demonstrated a proof-of-concept for the therapeutic targeting of P-cadherin, MCT1, or MCT4 using nanoparticle delivery. No mice exhibited apparent signs of toxicity, such as decreased food intake or activity or body weight loss, upon AmDM-siRNA treatment (data not shown). Thus, peritoneal dissemination may offer a clinical advantage in that therapeutic genes may be sensitive and specific to target cells owing to the closed space of the peritoneal cavity.

In conclusion, this study has uncovered a novel mechanism by which metastatic cells utilize adhesion-driven mechanical cues to shape their metastatic niche and promote metastatic outgrowth. Our findings emphasize the crucial role of P-cadherin in connecting mechanotransduction and cell metabolism within the peritoneal metastatic niche. Targeting the unique mechanoresponsive metabolic vulnerabilities of cancer cells and the tumor microenvironment holds promise for precise and localized therapeutic interventions, reducing side effects and overcoming treatment resistance. These findings may extend beyond ovarian cancer, as P-cadherin is also upregulated in other cancer types like gastric, pancreatic, and colon cancers, where peritoneal metastasis represents a significant pathological process.

## Methods

### Cell isolation and culture

HEYA8 HM and NM ovarian cancer cell sublines were constructed and maintained as previously described^18,19^. In brief, isogenic HM (highly metastatic) and NM (non-metastatic) cell lines were established by isolating migrated and non-migrated HEYA8 cell clones, respectively, from migration assay, and the differential metastatic properites were confirmed by orthotopic injection. The ovarian cancer cell line OVSAHO and the immortalized human mesothelial cells MeT5A cells were obtained from JCRB and ATCC respectively. Primary human peritoneal mesothelial cells (HPMC) were isolated from dialysate effluent from peritoneal dialysis from patients with nonmalignant disorders as previously described^58^. The collection of PD fluid was approved by the Institutional Review Board of the University of Hong Kong/Hospital Authority Hong Kong West Cluster. HPMC were used within three passages to ensure genetic stability of the culture. Normal human ovarian surface epithelial (OSE) cells were derived from surface scrapings of normal ovaries from women with nonmalignant gynecological diseases as previously described^59^. MeT5A, OSE and HPMC were all maintained in Medium 199:MCDB105 (1:1) supplemented with 100 U/mL penicillin and 100 μg/mL streptomycin (Invitrogen).

### Transfection and inhibitor treatment

Transient transfection with siRNA oligos (Dharmacon) or overexpression plasmids were performed using siLentFect for RNAi (Bio-Rad) or Lipofectamine 2000 (Thermo Fisher Scientific), respectively. To establish cell lines with stable knockdown, cells were transduced with MISSION® shRNA Lentiviral Transduction Particles (Sigma) according to the manufacturer’s protocols. For MCT inhibition, cells were treated with 200 nM monocarboxylate transporter inhibitor AR-C155858 (MedChemExpress) for 24 h. For ACC1 inhibition, cells were pretreated with TOFA (Abcam, 20 μM) for 2 h and then cocultured with MeT5A monolayer for 24 h.

### Cell adhesion assay

Mesothelial cells were seeded on 96-well plates and cultured to form confluent monolayer. CMFDA (Invitrogen)-labeled cancer cells were then added to the monolayer at a concentration of 1×10^4^ cells per well and incubated at 37℃ for 30 min. After gentle washing for three times with PBS, the plates were loaded on Cytation^TM^ 1 (BioTek) for cell counting at the GFP channel. The experiment was carried out in triplicate and repeated three times independently.

### Single-cell force spectroscopy

Cancer-mesothelial cell rupture forces were measured using JPK NanoWizard II (Bio-AFM). MeT5A cells were seeded at a density of 10^4^ cells/well on the 30 mm glass bottom dish (MatTek) and cultured overnight. Cantilevers were coated with 0.1 μg mL^−1^ poly-D-lysine solution (Sigma) for 6 h, rinsed with water before calibration using the thermal noise method. For cell-cell rupture force measurement, single cancer cell was picked up onto the cantilever with contact for 5 s, after which cancer cell was brought to contact with single MeT5A cell (medium changed to FBS-free RPMI 1640) for 5 s, with approach/retraction velocities of 5 μm s^−1^ and contact force of 2 nN. Force curves were collected and analyzed with JPK data processing software. For P-cadherin blocking group, HM and MeT5A cells were pre-incubated with 10 μg mL^−1^ P-cadherin antibody (NCC-CAD-299) (Thermo Fisher Scientific) for 15 min, after which rupture forces were measured as stated above.

### Western blot analysis

After SDS-PAGE, proteins were transferred to nitrocellulose membranes, which were blocked with 5% non-fat dry milk for 1 hand incubated with primary antibodies for 1 h to overnight. The membranes were washed thrice and incubated with appropriate secondary antibodies conjugated with HRP (Bio-rad; 1:3,000). Bands were detected by Western-Lightning Plus Enhanced Chemiluminescence (Perkin Elmer), and their intensities were determined by densitometry using ImageJ. P-cadherin (BD Biosciences, #610227), E-cadherin (CST, #3195), N-cadherin (CST, #13116), Cadherin-6 (CST, #48111), Cadherin-17 (CST, #85724), FASN (CST, #3180), Lipin 1 (CST, #14906), SREBP1 (Santa Cruz, #sc-13551), and β-actin (Sigma, #A5316) were used.

### Immunofluorescence microscopy

Cells were fixed with ice-cold methanol for 20 min and permeabilized with 1% Triton X-100 for 30 min. Cells were washed with PBS and blocked with 5% BSA in PBS for 20 min, followed by incubation with primary antibody (1:100) overnight. After washing with PBS, cells were incubated with dye conjugated secondary antibody (1:100) for 1 h. Stained cells were mounted with Prolong antifade mountant with DAPI (Invitrogen).

### Oil red O (ORO) staining

ORO staining was carried out using Lipid Staining Kit (BioVison). Briefly, cells were treated as indicated and fixed with Formalin (10%) for 30 min before staining with ORO in 60% isopropanol for 15 min. Confocal imaging was used to capture the Oil red O signals. Only signals overlaying with green fluorescence (from tumor cells labeled with CellTracker^TM^ Green) were measured.

### Glucose uptake assay

Cells were inoculated in a 6-well plate in 5% FBS supplemented culture medium. After the indicated treatment for 24 h, medium was changed to glucose-free medium with 100 μg/mL 2-NBDG. Cells were incubated for 45 min and washed twice with ice cold PBS, before loading on Cytation^TM^ 1. More than 3 technical replicates were analyzed for cell counting by ImageJ.

### L-Lactate measurement

The L-lactate levels of cells or culture medium were measured by L-Lactate Assay Kit (Abcam). Cells were collected by centrifugation and resuspended in 100 μL lactate assay buffer, which were then homogenized and centrifuged to collect supernatant for L-lactate assay. The culture medium was measured directly according to the manufacturer’s instructions.

### Seahorse extracellular flux analysis

Mitochondrial and glycolysis bioenergetics were investigated using Seahorse Extracellular Flux analyzer (Agilent). Cells were treated with siRNA for 24 h and reseeded onto 24-well XF plates for another 24 h before bioenergetics analysis. Growth medium was then changed with XF base medium (pH, 7.4) containing 25 mM glucose, 1 mM sodium pyruvate, and 2 mM glutamine for mitochondrial bioenergetics assay, or 2 mM glutamine (Sigma) for glycolysis bioenergetics assay. Oxygen consumption rate (OCR) or extracellular acidic rate (ECAR) profiles were generated after the injection of 1.5 μM oligomycin, 3 μM FCCP and 1.2 μM rotenone+1.2 μM antimycin A for OCR, or 15 mM Glucose, 1.5 μM oligomycin, 75 mM 2-DG for ECAR every 24 min. The average of four baseline rates and up to five test rates were used for data analysis.

### Proliferation assay

HM cells labeled with CMFDA were inoculated on the mesothelial monolayer or alone in RPMI 1640 supplemented with 5% FBS, and monitored every 24 h for 4 days. More than 3 technical replicates were analyzed for cell counting by ImageJ.

### ^13^C6-glucose and lactate competition assay

The lactate competition assay was modified from reported method^60^. Briefly, mesothelial monolayer was first placed in RPMI 1640 (Gibco) containing 25 mM [U^13^C]-glucose (Cambridge Isotope Laboratories) for 24 h. The monolayer was then washed with glucose-free RPMI 1640 and labeled with CellTracker^TM^ Blue CMAC Dye (Invitrogen) for 45 min before coculture with CellTracker^TM^ Green CMFDA Dye (Invitrogen)-labeled HM cells. The cells were further cultured in RPMI-1640 supplemented with 25 mM [U^13^C]-glucose in the absence or presence of 11 mM L-lactate (Sigma).

### ^13^C-metabolic flux analysis

^13^C-lactate tracing was performed using female mice which were first intraperitoneally (i.p). injected with NM or HM cells. After 3 weeks, during which the metastatic tumors had formed, mice were fasted for 3 h before receiving i.p. injection of 10 mM ^13^C_3_ sodium lactate (Cambridge Isotope Laboratories). After 3 h, mice were sacrificed and the metastatic tumors were harvested for targeted lipidomics analysis. For ^13^C-glucose flux analysis, MeT5A seeded in 100 mm dish were cultured in glucose-free medium containing 25 mM [U^13^C_6_]-glucose for 24 h. CellTracker^TM^ Green CMFDA Dye (Invitrogen)-labeled HM cells transfected with siP-cad or siNS were subsequently added for coculture. After 24 h, cells were collected and HM cells were isolated by fluorescence-activated cell sorting. Cells were extracted and subjected to GC-MS/MS analysis through an Agilent DB-23 capillary column and acquired in an Agilent 7890B GC-Agilent 7010 Triple Quadrapole Mass Spectrometer system (Santa Clara, USA). For fatty acids flux analysis, fatty acids were determined after transesterification by methanol and concentrated hydrochloric acid (35%, w/w). For determination of PEP and pyruvate, extracted samples by methanol/water (80%, v/v) were derivatized with methoxylamine hydrochloride and MSTFA with 1% TMCS. Metabolite fragment ions of monoisotopic mass (denoted M + 0) and mass isotopologues of higher mass were quantified in SIM mode. A scan range from m/z 50-500 was used to acquire mass spectra in SCAN mode. Data analysis was performed using the Agilent MassHunter Workstation Quantitative Analysis Software. Relative fractions of mass isotopologues were calculated from the peak areas of corresponding SIM ions. Raw data acquisition was performed with assistance of the University of Hong Kong Li Ka Shing Faculty of Medicine Centre for PanorOmic Sciences Proteomics and Metabolomics Core.

### Cell sorting

After cocultured for 24 h, fluorescent-labeled cells were sorted by BD FACSMelody^TM^ cell sorter, and collected in RPMI 1640 supplemented with 5% FBS at 4℃.

### Treatment with Fc- or P-cadherin/Fc-coated beads

Magnetic beads were coated with recombinant human P-cadherin/Fc chimera protein or recombinant human IgG1 Fc protein (R&D systems) according to manufacturer’s instructions. Briefly, 1.5 mg Dynabeads Protein A (Invitrogen) were incubated with 50 μL Fc or P-cad/Fc protein (100 μg/mL) in 200 μL PBS (0.1% Tween-20) for 10 min at room temperature. Beads were collected using magnet to remove unbound protein and washed once with PBS with 0.1% Tween-20. The coated beads were stored at 4 ℃ and used within one week.

### RNA extraction and reverse-transcription PCR analysis

Total RNA was first extracted using Trizol reagent (Ambion) and then reverse-transcribed to cDNA using M-MLV reverse transcriptase (Invitrogen). PCR was performed with a set of primers as listed in Supplementary Table 2.

### RNA sequencing and data analysis

RNA was extracted from HM cells were transfected with NS or P-cadherin siRNA and sent to BGI for transcriptomic RNA sequencing (n=2). Data have been deposited in the Genome Sequence Archive (China National Center for Bioinformation) which are publicly accessible at https://ngdc.cncb.ac.cn/gsa-human (GSA-Human: HRA004687). Dysregulated pathways were identified with Generally Applicable Gene-set Enrichment (GAGE).

### siRNA dendriplexes formation

AmDM (MW=3838 g/mol, 16 amine end groups) were dissolved in distilled H_2_O and stocked in 500 μM. To prepare the siRNA dendriplexes, dendrimer and siRNA were first diluted separately in OPTI-MEM (Invitrogen) and incubated at room temperature for 5 min. Dendrimer and siRNA were then mixed at different N/P ratios (the number of amino end group on dendrimer to the number of phosphate end group on siRNA) or different siRNA concentrations and incubated at room temperature for 20 min before treatment.

### In vivo study

All mouse studies were performed according to the protocols approved by the University of Hong Kong Animal Care and Use Committee. Female NOD/SCID/IL2rγ^null^ (NSG) mice were purchased from Charles River Laboratories. HM, NM or transfected HM cells (1 × 10^6^) were intraperitoneally injected into NSG mice (n=3 mice per group) and sacrificed after 3 weeks. For dendriplexes treatment, mice bearing intraperitoneally injected HM cells were intravenously injected with siNS-AmDM, siP-cad-AmDM, siMCT1-AmDM or siMCT4-AmDM for 3 weeks (siRNA: 1.0 mg/kg, N/P ratio=5, twice a week). At sacrifice, the volume of ascites was measured and all visible (> 0.1 cm) metastatic tumor nodules were counted.

### Human patient samples

Formalin-fixed paraffin-embedded (FFPE) clinical samples of paired ovarian high-grade serous carcinoma and metastases were obtained from the pathology archives of Department of Pathology, University of Hong Kong, for immunohistochemical analyses. Clinicopathological information of patients were provided in Supplementary Table 1. Primary tumor ascitic samples were obtained from 7 high grade serous ovarian cancer patients (Stage III/IV). The use of these specimens was approved by the Institutional Ethical Review Board for Research on the use of human subjects at the University of Hong Kong.

### Immunohistochemistry

FFPE sections from patient or mice tumors were deparaffinized with xylene and rehydrated in graded ethanol. After blocking and heat-induced antigen retrieval using citrate buffer, the specimens were separately incubated with primary antibody (FASN (#3180, CST, 1;100), P-cadherin (ab242060, Abcam, 1:1000), Ki67 (#9027, CST, 1:1000)) overnight at 4 ℃. The slides were further processed using rabbit specific HRP/DAB (ABC) detection IHC kit according to the manufacturer’s protocol (Abcam). Slides were subsequently scanned on an Akoya Vectra Polaris scanner scanning at ×20 magnification and semi-quantitative analysis of cytoplasmic FASN, membrane P-cadherin and nuclear Ki-67 expression was performed using QuPath 0.4.4 software. Five fields per specimen with the strongest marker expression were selected for cell detection and tumor cell classification. Followed by single cell detection, staining intensity (0, negative; 1, weak; 2, medium and 3, strong) and percentage of positive tumor cells were assessed based on pre-defined arbitrary signal intensity. Gene expression was graded with an H-score as previously reported, which is calculated by multiplying the percentage of positive cells (0-100) by the intensity (0-3). Data are expressed as median H-score.

### Statistical analyses

All experiments were carried out in duplicates or triplicates and repeated at least two times. All data are presented as mean ± SD unless otherwise indicated. Data derived from two groups were compared by unpaired Student’s *t* test. Data derived from more than two groups were compared by one-way ANOVA followed by a Tukey’s test. Paired sample *t* test is used for comparing expression of markers in paired patient specimens. Differences were considered statistically significant when the *P* value was less than 0.05.

## Data availability statement

The authors confirm that the data supporting the findings of this study are available within the article and its supplementary materials. RNA-seq data have been deposited in the Genome Sequence Archive (China National Center for Bioinformation) (GSA-Human: HRA004687) which are publicly accessible.

## Acknowledgements

This work was supported by the Research Grant Council General Research Fund 17105919 and Senior Research Fellow Scheme SRFS2223-7S05 to A. S. T. Wong, and the funding support from “Laboratory for Synthetic Chemistry and Chemical Biology” under the Health@InnoHK Program launched by Innovation and Technology Commission, HKSAR. This work was supported by the Hong Kong Scholars Program [XJ2019051].

## Ethics approval statement

All mouse studies were performed according to the protocols approved by the University of Hong Kong Animal Care and Use Committee. The use of patient specimens was approved by the Institutional Ethical Review Board for Research on the use of human subjects at the University of Hong Kong.

## Author contributions

A.S.T.W., J.M., S.KY.T., and K.W. designed the research and prepared the manuscript. J.M., S.K.Y.T., K.S.W.F., and K.W. performed research and analyzed data. J.Z. performed the bioinformatics analysis. S.Y. and T.C. provided primary human peritoneal mesothelial cells. C.C.L.W. assisted lactate assay. P.P.C.I. provided clinical samples. L.P. provided dendrimers. C.B.C., A.H.W.N. and H.Y.G. provided key knowledge. C.B.C. and A.S.T.W. supervised the study.

