## Supplementary Figures for "P-cadherin mechanoactivates tumor–mesothelium metabolic coupling to promote ovarian cancer metastasis"

**Supplementary Fig. 1**

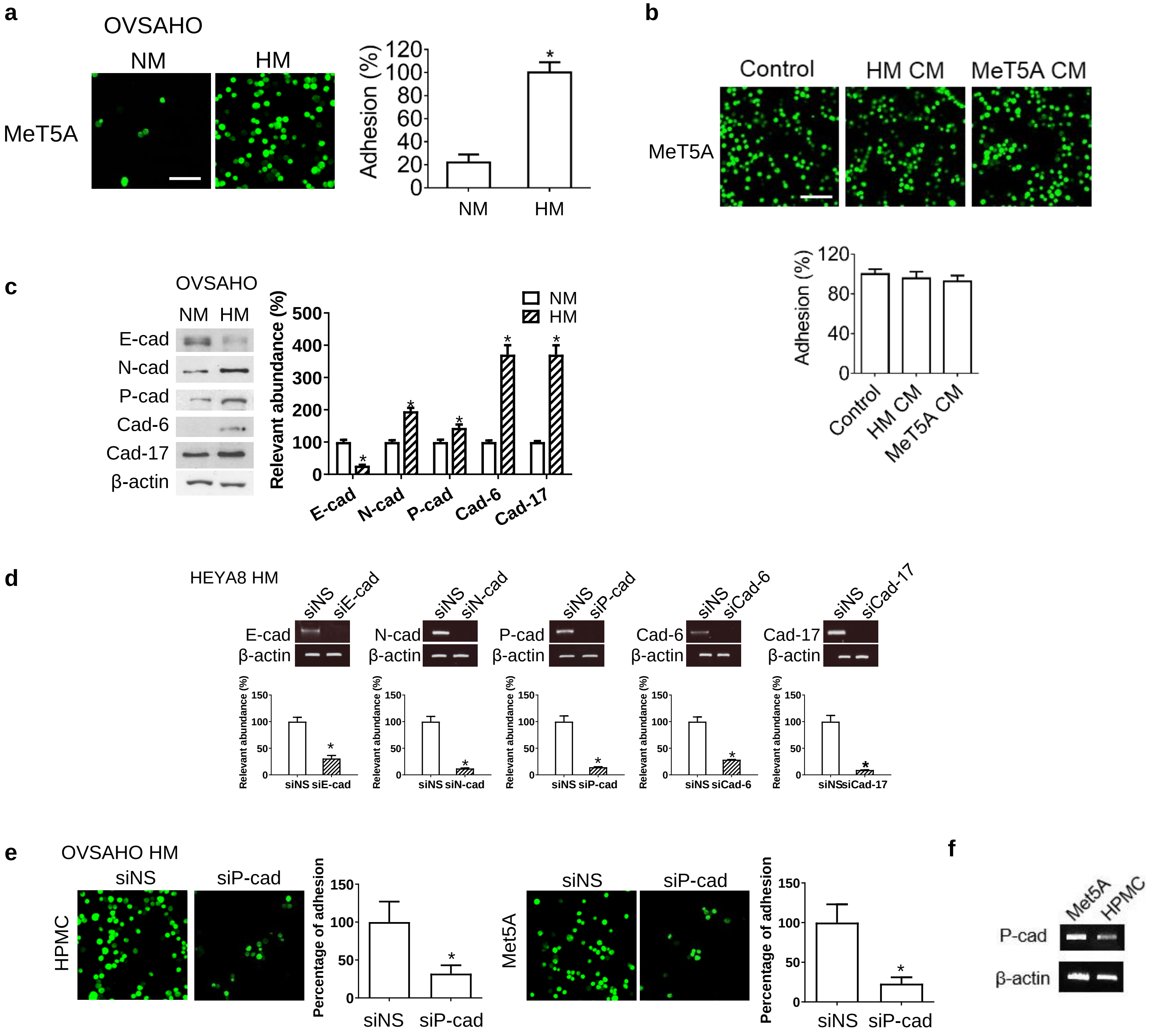

**Supplementary Fig. 1 a,** Representative fluorescent images and quantification of OVSAHO NM or HM cells (labelled with CMFDA, green) adhered onto a confluent MeT5A monolayer are shown. Scale bar, 100 µm. **b**, Representative fluorescent images and quantification of adhesion percentage of HEYA8 HM (green) to MeT5A monolayer in the absence or presence of conditioned medium (CM) as indicated. Scale bar, 100 µm. **c,** Expression of E-cadherin (E-cad), N-cadherin (N-cad), P-cadherin (P-cad), cadherin-6 (Cad-6), and cadherin-17 (Cad-17) in OVASHO HM and NM cells were analyzed by Western blot, with β-actin as the loading control. *$P < 0.05$ vs. NM. **d,** RT-PCR analysis and quantification to validate siRNA knockdown of E-cad, N-cad, P-cad, Cad-6 and Cad-17 in HEYA8 HM cells. Data are presented as mean percentage of knockdown ± SD. **e,** Representative images and quantification of adhesion percentage of siNS or siP-cad treated OVSAHO HM cells (green) adhered on primary human peritoneal mesothelial cell (HPMC) and MeT5A monolayer. **f,** The expression of P-cadherin in Met5A and HPMC are measured by RT-PCR analysis. Data are presented as mean ± SD. *$P < 0.05$.

### Supplementary Fig. 2

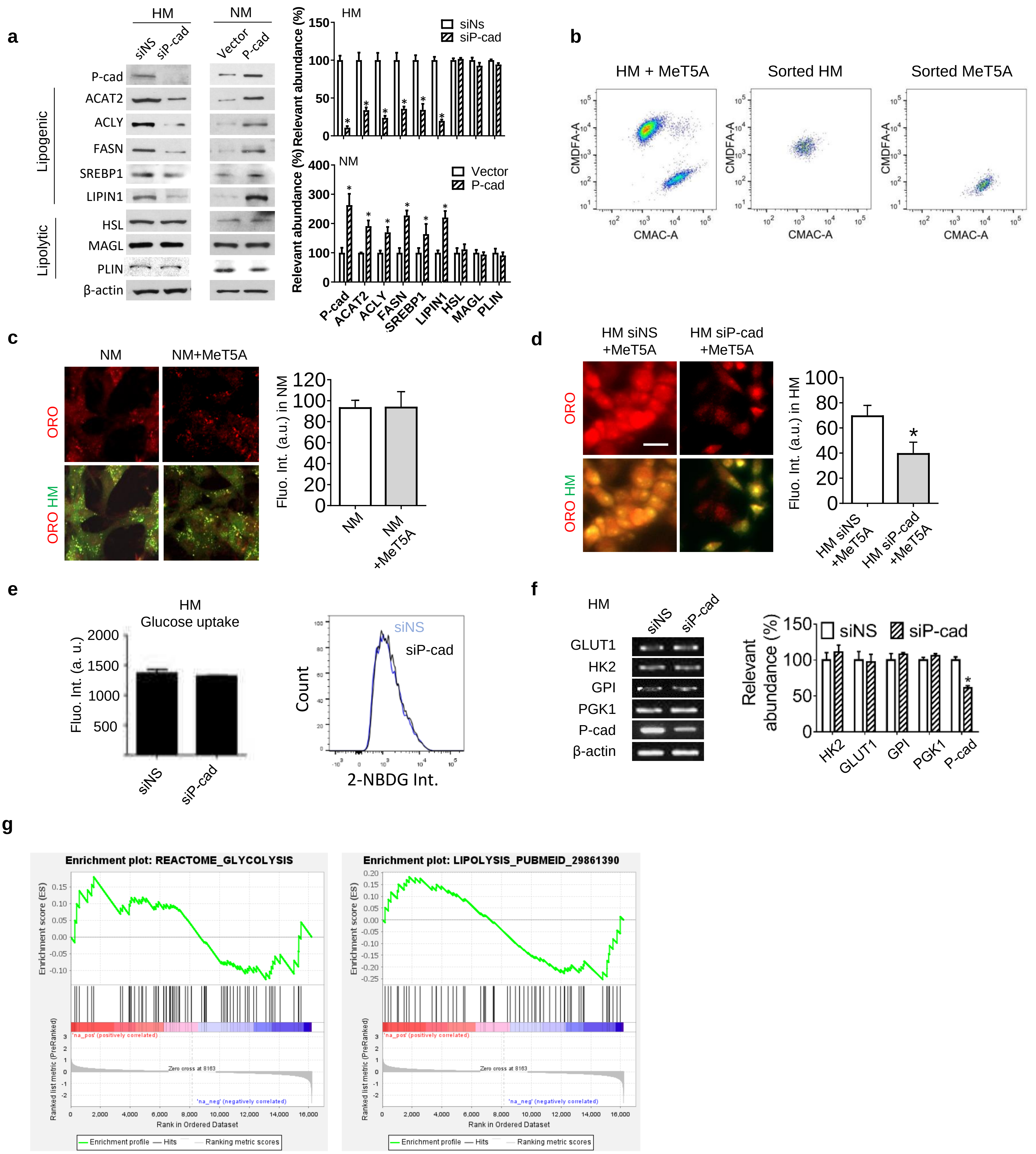

**Supplementary Fig. 2 a,** Western blot analysis of the expression of P-cadherin (P-cad), lipogenic genes (ACAT2, ACLY, FASN, SREBP1 and LIPIN1) and lipolytic genes (HSL, MAGL and PLIN) in HM or NM cells transfected with siNS, siP-cad, empty vector or P-cad overexpression vector as indicated. **b,** Representative flow cytometry profiles of HM and MeT5A cells (stained by CMDFA and CMAC, respectively) before and after flow cytometric sorting. **c-d,** Representative images and quantification of oil red O (ORO) staining of neutral lipids (red) in CMFDA-labelled (green) **(c)** NM cells with or without MeT5A coculture and **(d)** HM cells transfected with siNS or siP-cad before MeT5A coculture. Scale bar, 20 μm. **e,** Flow cytometry analysis and quantification of glucose uptake in siNS or siP-cad treated HM cells using 2-NBDG. **f,** RT-PCR analysis and quantification of glycolytic genes (GLUT1, HK2, GPI and PGK1) and P-cad in HM cells transfected with siNS or siP-cad. β-actin was used as the loading control. **g,** GSEA enrichment plot of glycolytic and lipolysis genes in HM cells in comparison to NM cells. Glycolysis gene set is from REACTOME_GLYCOLYSIS. Lipolysis gene set is from the https://doi.org/10.1016/j.cmet.2018.05.004 (lipolysis genes (n = 67). Data are expressed as mean ± SD. *$P < 0.05$.

### Supplementary Fig. 3

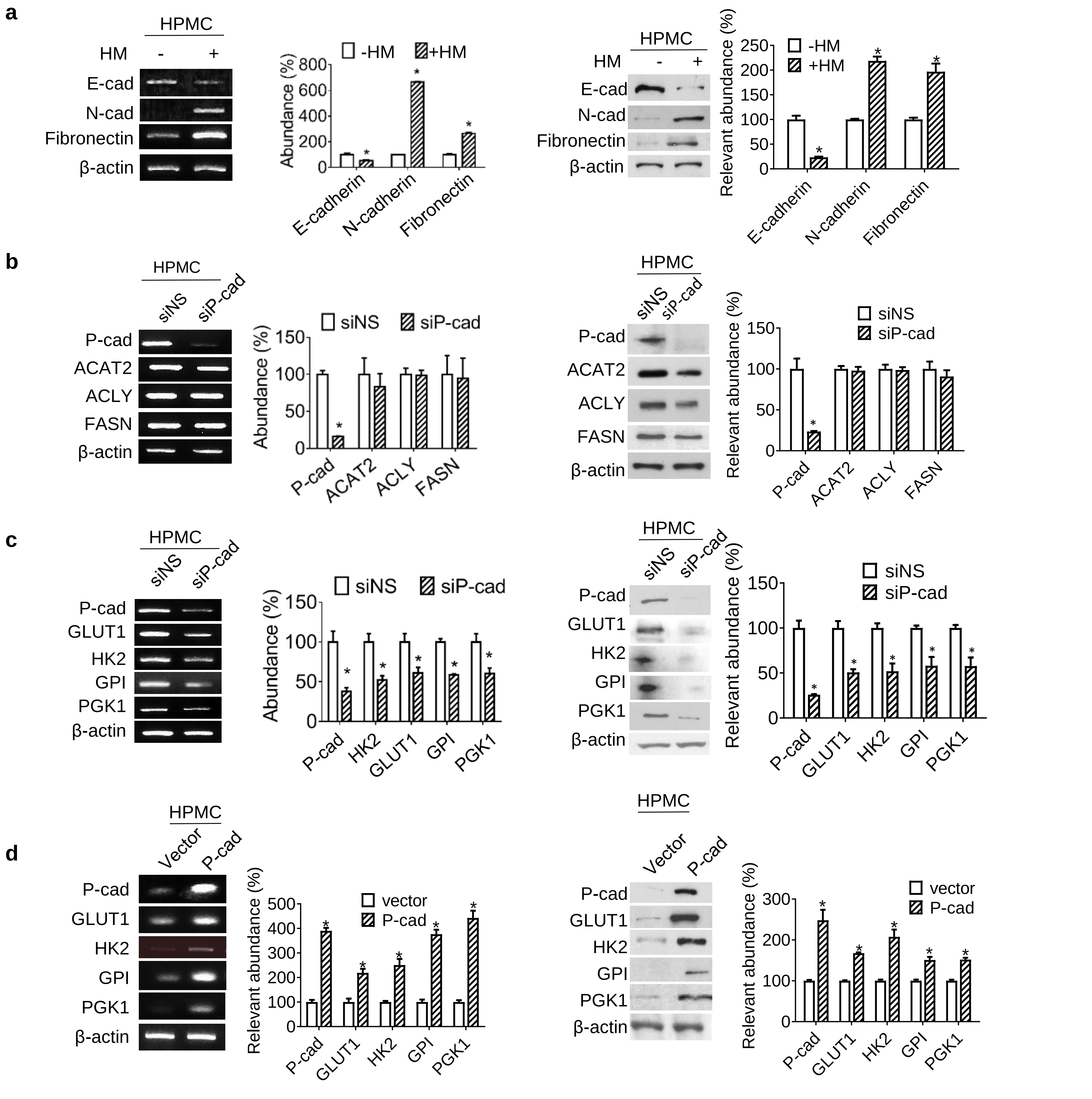

**Supplementary Fig. 3 a,** RT-PCR analysis and Western blot analysis of E-cadherin (E-cad), N-cadherin (N-cad), and fibronectin levels in primary human peritoneal mesothelial cells (HPMC) cocultured with or without HM cells. **b,** RT-PCR analysis and Western blot analysis of P-cadherin (P-cad) and lipogenic genes (ACLY, FASN and ACAT2) in siNS or siP-cad treated HPMC. **c,** RT-PCR analysis and Western blot analysis of P-cad and glycolytic genes (GLUT1, HK2, GPI and PGK1) in siNS or siP-cad treated HPMC cells. **d,** HPMC transfected with P-cad overexpression or empty vector were cocultured with HM. After coculture, HPMC were sorted by flow cytometry, followed by RT-PCR and Western blot analysis of P-cad and glycolytic genes (GLUT1, HK2, GPI, and PGK1). β-actin serves as the loading control for all RT-PCR and Western blot analysis. Quantification analysis were performed, and data are presented as mean ± SD. *$P < 0.05$.

**Supplementary Fig. 4**

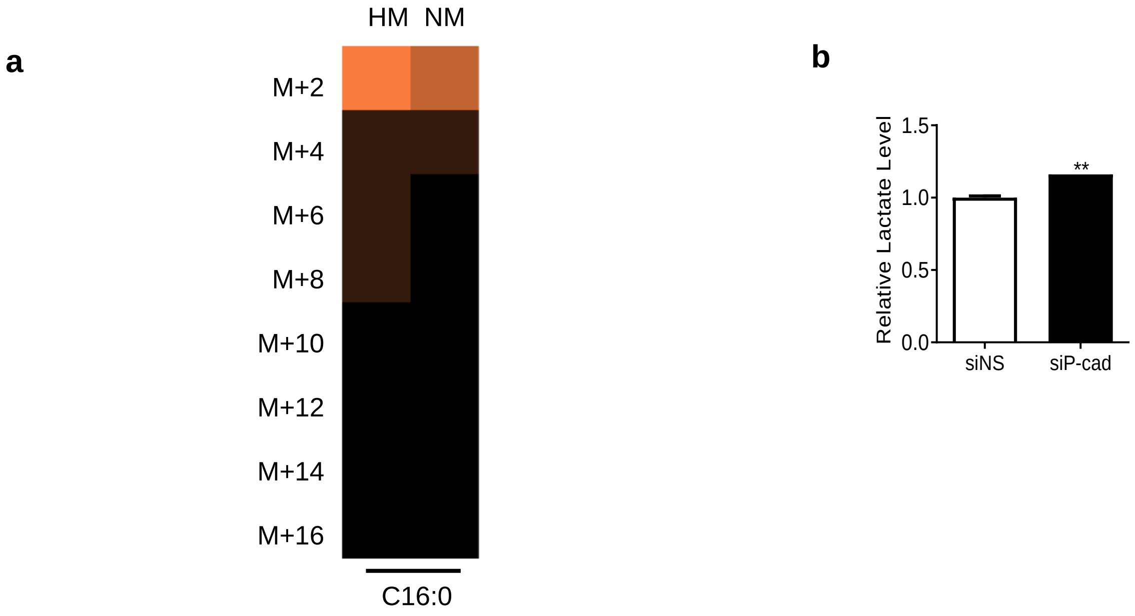

**Supplementary Fig. 4 a,** Lipidomic analysis of $^{13}C$-labeled palmitic acid (C16:0) in HM or NM tumor nodules *in vivo.* **b,** HM cells transfected with siNS or siP-cad were treated with P-cad/Fc-coated beads. Extracellular lactate levels in the conditioned media were measured.

### Supplementary Fig. 5

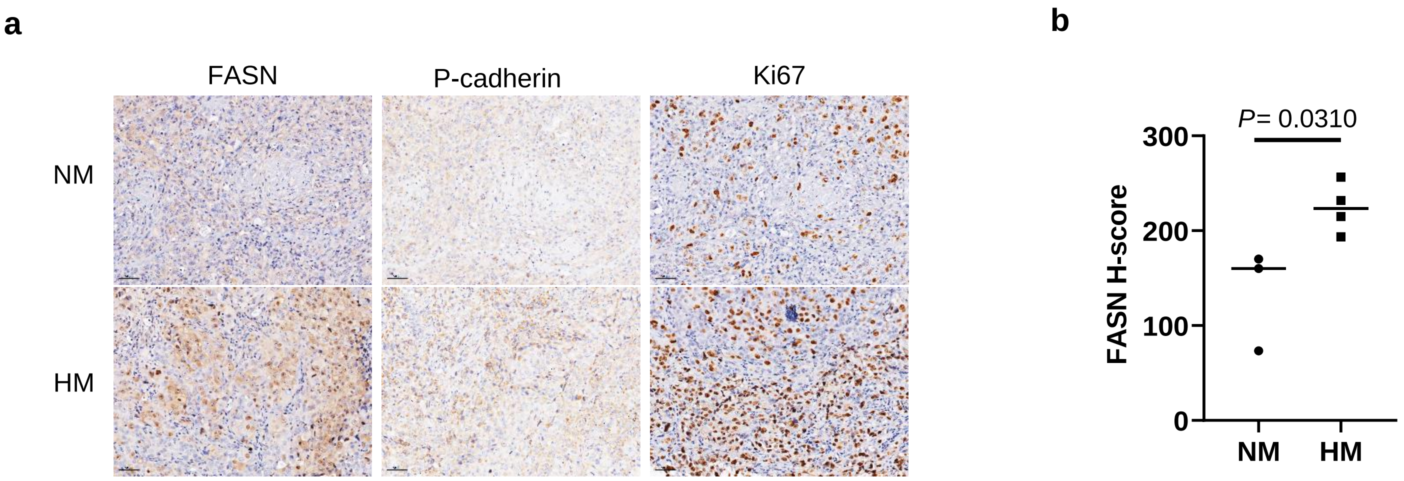

**Supplementary Fig. 5 a,** Representative images of immunohistochemical analysis of FASN, P-cadherin and Ki67 in NM and HM tumors in mice. **b,** FASN expression was quantified in NM (n=3) and HM tumors (n=4) using H-score.

Supplementary Fig. 6

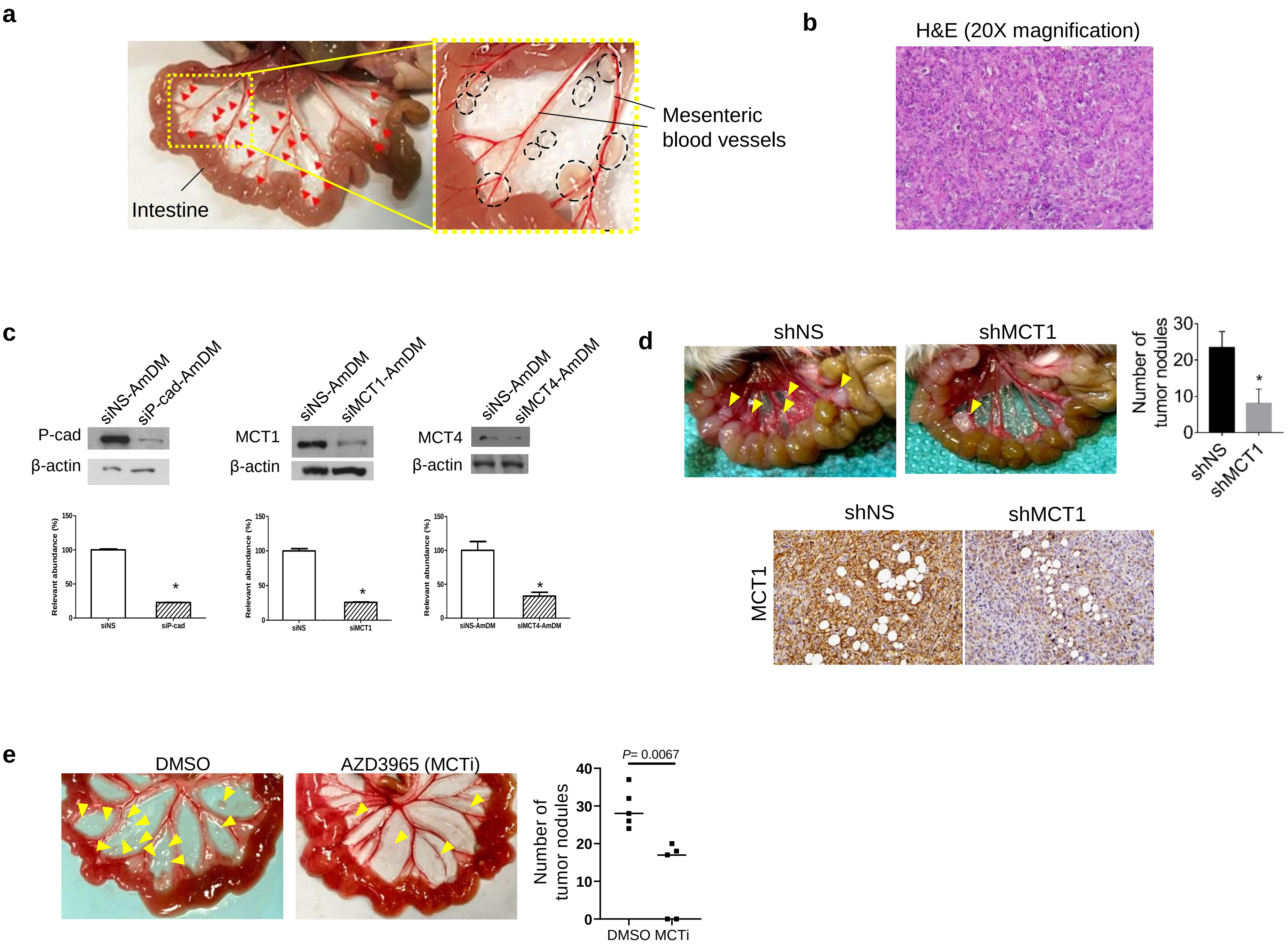

**Supplementary Fig. 6 a,** Representative images showing the gross anatomy of mesentery with peritoneal metastatic nodules. **b,** Hematoxylin and eosin (H&E) staining showing histological phenotype of HM tumors. **c,** Western blot analysis shows the knockdown of P-cad, MCT1 and MCT4 in HM cells after treatment of siP-cad-AmDM, siMCT1-AmDM and siMCT4-AmDM, respectively. *$P$ < 0.05 compared to siNS-AmDM. **d,** HM cells treated with non-specific or MCT1 shRNA (shNS and shMCT1) were intraperitoneally injected into mice. Representative images and the number of metastatic nodules were shown (upper panel). Immunohistochemical images shows the expression of MCT1 in shNS and shMCT1 tumors (lower panel). **e,** Mice with intraperitoneally injected HM cells are treated with DMSO (n= 5) or monocarboxylate transporter inhibitor (MCTi, AZD3965) (n= 5). Representative images and the number of metastatic nodules were shown.

**Supplementary Table 1. Summary of clinical characteristics of paired primary and metastatic samples from patients.**

| | **Metastatic specimens (n=16)** |
|---|---|
| Characteristics | |
| High grade serous ovarian carcinoma | 16 |
| FIGO stage | |
| IIIB | 1 |
| IIIC | 11 |
| IV | 4 |
| Age (range 35-78) | |
| <60 | 12 |
| ≥60 | 4 |
| Site of metastatic biopsy | |
| Omentum | 9 |
| Bladder flap | 4 |
| Pelvic peritoneum/ peritoneal | 3 |

**Supplementary Table 2. Primer sequences used for RT-PCR assay.**

| Genes | 5' Primer | 3' Primer |
|---|---|---|
| ACAT2 | CCTGTGGTCATCGTCTCGGC | AAGCCAAGTGAGGAGCCTTG |
| ACLY | AAGGGCATCGTGAGAGCAAT | TGGGACTGAATCTTGTGGCAT |
| β-actin | TCACCGAGGCCCCTCTGAACCCTA | GGCAGTAATCTCCTTCTGCATCCT |
| CDH1 | ACAGCCCCGCCTTATGATT | TCGGACCGCTTCCTTCA |
| CDH2 | TGGGAATCCGACGAATGG | GCAGATCGGACCGGATACTG |
| CDH3 | ACGAAGACACAAGAGAGATTGG | AGCAACCACCCCATTGTAGG |
| CDH6 | TCAAGACAACAAAGACAACACG | TCTCCACCACCTTCGTCGTT |
| CDH17 | ATGACAACCCTCCCAGGGC | GACCCCGAAATAAGTGCTGA |
| FASN | CTGGCCTACACCCAGAGCTA | CTCCATGTCCGTGAACTGCT |
| Fibronectin | AAGACAGACGAGCTTCCCCA | CTATGCCTTATGGGGGTGGC |
| Gpi | GCGAGGCCAGAGTCCAATAA | GACCAGCCTTCCCTGCTAAG |
| GLUT1 | GCAAGTCCTTTGAGATGCTGATCC | GCCGACTCTCTTCCTTCATATCC |
| HK2 | CCAGTTCATTCACATCATCAG | CTTACACGAGGTCACATAGC |
| HSL | TCAGTGTGCTCTCCAAGTGTGTC | GCTATGGGCTCCGACATCTTC |
| MAGL | TTGCTGCGAAAGTGCTCAAC | ATTCAGCAGTTGGATGCCGA |
| Perilipin1 | GAAACAGCATCAGCGTTCCC | ATGGTCTGCACGGTGTATCG |
| PGK1 | CTGTGCCAAATGGAACACGG | AGTTGACTTAGGGGCTGTGC |
| SLC16A1 | TGGATGGAGAGGAAGCTTTCTAAT | CACACCAGATTTTCCAGCTTTC |
| SREBP1 | ACAGTGACTTCCCTGGCCTAT | GCATGGACGGGTACATCTTCAA |
